# Insights into stereoselective ring formation in canonical strigolactone: Discovery of a dirigent domain-containing enzyme catalyzing orobanchol synthesis

**DOI:** 10.1101/2023.08.07.552212

**Authors:** Masato Homma, Takatoshi Wakabayashi, Yoshitaka Moriwaki, Nanami Shiotani, Takumi Shigeta, Kazuki Isobe, Atsushi Okazawa, Daisaku Ohta, Tohru Terada, Kentaro Shimizu, Masaharu Mizutani, Hirosato Takikawa, Yukihiro Sugimoto

## Abstract

Strigolactones (SLs) are plant apocarotenoids with diverse functions and structures. The widespread canonical SLs, with distinctive structural variations in their tricyclic lactone known as the ABC-ring, are classified into two types based on the C-ring configurations. The steric C-ring configuration arises during the BC-ring closure downstream of carlactonoic acid (CLA), a biosynthetic intermediate. Most plants stereoselectively produce either type of canonical SLs, e.g., tomato (*Solanum lycopersicum*) produces orobanchol with α-oriented C-ring. The mechanisms governing SL structural diversification are partly understood, with limited insight into the functional implications. Moreover, the precise molecular mechanism for the stereoselective BC-ring closure reaction remains unknown. Herein, we identified an enzyme called the stereoselective BC-ring-forming factor (SRF) from the dirigent protein (DIR) family, especially the DIR-f subfamily, whose biochemical function was previously unidentified, making it a pivotal enzyme in stereoselective canonical SL biosynthesis with the α-oriented C-ring. We begin by confirming the exact catalytic function of the tomato cytochrome P450 SlCYP722C, which we previously demonstrated to be involved in the orobanchol biosynthesis [Wakabayashi et al., *Sci. Adv.* **5**, eaax9067 (2019)], to convert CLA to 18-oxocarlactonoic acid. Subsequently, we demonstrate that SRF catalyzes the stereoselective BC-ring closure reaction of 18-oxocarlactonoic acid to form orobanchol. Our approach integrates experimental and computational methods, including SRF structure prediction and molecular dynamics simulations, to propose a catalytic mechanism based on the conrotatory 4π-electrocyclic reaction for stereoselective BC-ring formation in orobanchol. The present study provides insight into the molecular basis of how plants produce SLs with specific stereochemistry in a controlled manner.

## Introduction

Strigolactones (SLs) comprise a group of plant apocarotenoids with various functions^1^. SLs are endogenous plant hormones that inhibit shoot branching/tillering^2, 3^, rhizosphere signaling molecules secreted from roots for hyphal branching of arbuscular mycorrhizal fungi^4^, and germination stimulants for root parasitic weeds^5^. SLs are diverse and more than 30 structures have been reported thus far^6^. Plants share a common upstream biosynthesis pathway for SL, wherein all*-trans*-β*-*carotene is converted to the biosynthetic intermediate carlactone (CL) through the sequential catalysis by the three enzymes, carotenoid isomerase DWARF27 (D27), carotenoid cleavage dioxygenase 7 (CCD7), and CCD8. The cytochrome P450 CYP711A subfamily catalyzes the conversion of CL to carlactonoic acid (CLA), which is a conserved function of this enzyme subfamily (Fig. 1)^7–11^. The structural diversification in SLs is known to primarily occur downstream of CLA synthesis. The ABC-ring system, comprising the tricyclic lactone moiety, embodies the most distinctive SL structural differences. Canonical SLs, which contain this ring system, are classified into two subgroups based on their C-ring configuration, the orobanchol- and strigol-types. The orobanchol-type SLs have α-oriented C-rings, whereas the strigol-type SLs have β-oriented C-rings. Conversely, non-canonical SLs have incomplete ABC-ring systems (Fig. 1). The mechanisms underlying the structural diversification of SLs have only been partially elucidated, and the functional implications of this structural diversity are also poorly understood. Recent studies have disclosed the possibility that the structurally different SLs have respective roles. Reportedly, certain non-canonical SLs with methyl-esterified substructures of CLA were suggested to act as unidentified shoot branching inhibiting hormones^12^, whereas canonical SLs are important signaling molecules in the rhizosphere^13, 14^. Most plants stereoselectively produce either type of canonical SLs and the differences in the C-ring configuration have physiological significance. For example, administration of SLs with different C-ring configurations results in altered physiological and gene expression responses in *Arabidopsis thaliana*^15, 16^. In the rhizosphere, the C-ring stereochemistry affects the germination-stimulating activity of root parasitic weeds^17^ and the recruitment of bacterial communities as found among sorghum (*Sorghum bicolor*) cultivars with different SL profiles^18, 19^.

**Fig. 1.**
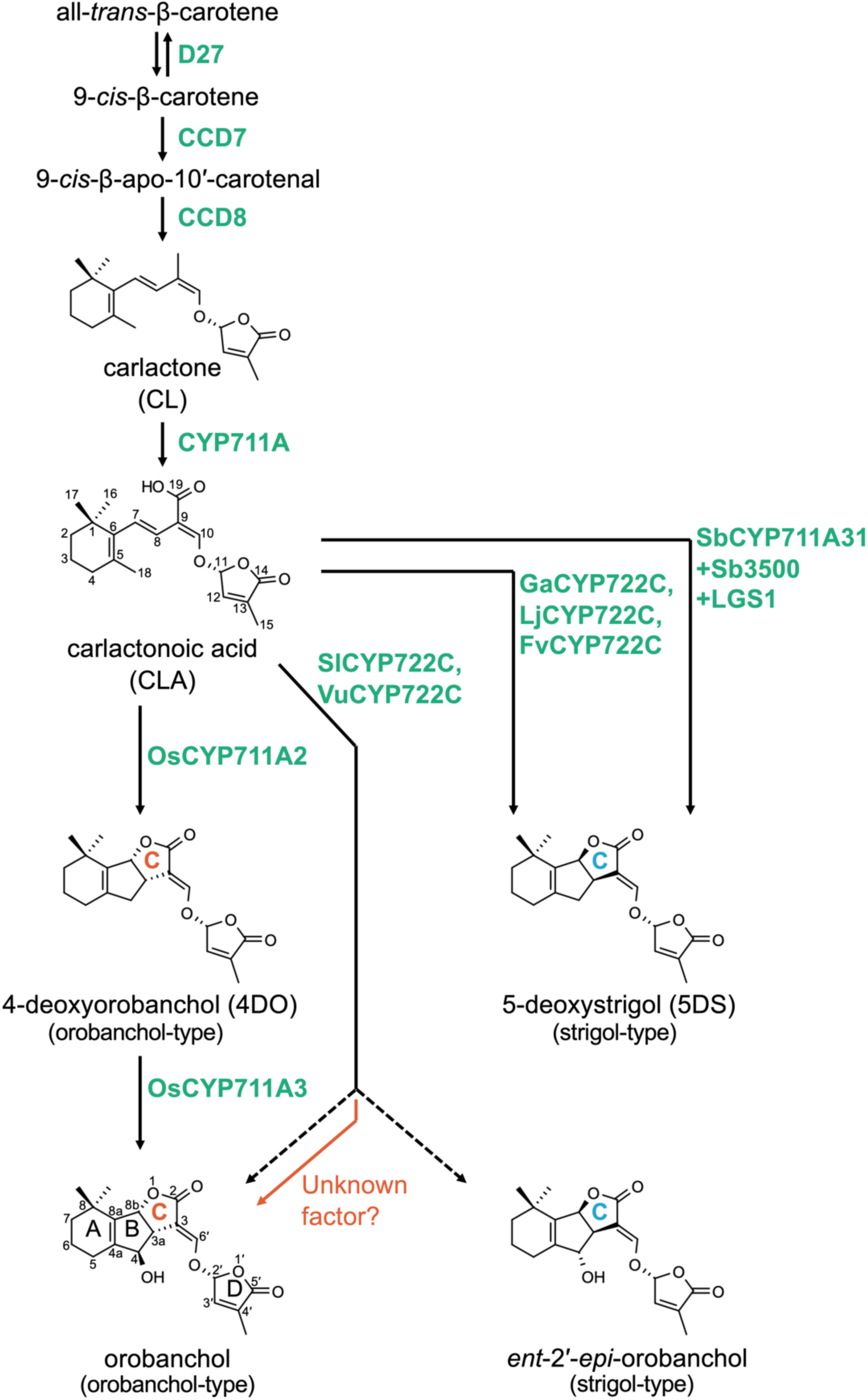
Representative biosynthesis pathway of canonical strigolactone (SL). The sequential reactions catalyzed by the D27, CCD7, and CCD8 enzymes produce carlactone (CL) from all- *trans*-β-carotene. CL is further oxidized by the CYP711A subfamily to produce carlactonoic acid (CLA). The types of enzymes involved in canonical SL biosynthesis differ depending on plant species. Several CYP711A subfamily members are implicated in the formation of canonical SL; OsCYP711A2 in rice is involved in the 4-deoxyorobanchol (4DO) biosynthesis, and in sorghum, SbCYP711A31 is suggested to function cooperatively with additional enzymes, Sb3500 and LGS1, to form 5-deoxystrigol (5DS). The CYP722C subfamily, which is widely conserved in dicots, is also involved in the formation of canonical SL from CLA. An additional factor which regulates the C-ring configuration was suggested to be present in the biosynthesis of orobanchol mediated by SlCYP722C and VuCYP722C in tomato and cowpea, respectively. Fv, *Fragaria vesca*; Ga, *Gossypium arboreum*; Lj, *Lotus japonicus*; Sb, *Sorghum bicolor*; Os, *Oryza sativa*; Sl, *Solanum lycopersicum*; Vu, *Vigna unguiculata*. LGS1, LOW GERMINATION STIMULANT 1.

The steric configuration of the C-ring of canonical SLs arises during the BC-ring closure reaction downstream of CLA (Fig. 1). OsCYP711A2/Os900 in rice (*Oryza sativa*) catalyzes the stereoselective BC-ring closure of CLA to form 4-deoxyorobancol (4DO), which is an orobanchol-type SL with α-oriented C-ring^10, 11^. In sorghum (*Sorghum bicolor*), it was just recently reported that the strigol-type SL 5-deoxystrigol (5DS) with β-oriented C-ring is stereoselectively generated from CL or CLA in vitro by the concerted catalytic action through an unknown reaction mechanism involving three different type enzymes, SbCYP711A31/SbMAX1a, a cytochrome P450, Sb3500, a 2-oxoglutarate-dependent dioxygenase, and LOW GERMINATION STIMULANT 1 (LGS1), a sulfotransferase^20^. The CYP722C subfamily, which is widely conserved in dicots, contains the key enzymes required for canonical SL synthesis. We previously demonstrated that GaCYP722C in cotton (*Gossypium arboreum*), which produces 5DS, catalyzes the conversion of CLA to 5DS^21^. The CYP722Cs in 5DS-producing plants, such as birdsfoot trefoil (*Lotus japonicus*) and woodland strawberry (*Fragaria vesca*), are also involved in this stereoselective conversion^22, 23^, which suggests a conserved function of the CYP722C subfamily in 5DS-producing dicot plants. Enzymes of this subfamily are also involved in the biosynthesis of the orobanchol-type SL orobanchol, which has been detected in the root exudates of numerous plants^24^. In contrast to the stereoselective synthesis of 5DS, however, *in vitro* enzyme assays using the substrate CLA for VuCYP722C and SlCYP722C from cowpea (*Vigna unguiculata*) and tomato (*Solanum lycopersicum*), which exclusively produce orobanchol and its related SLs, revealed that their enzyme products are the orobanchol diastereomers, orobanchol and *ent*-2′-*epi*-orobanchol that has the opposite C-ring configuration to that of orobanchol^13^ (Fig. 1). Knockout of the *SlCYP722C* gene in tomato resulted in the loss of orobanchol and the alternative accumulation of its substrate CLA, thereby suggesting that this enzyme is responsible for the initial step in the BC-ring closure reaction^13^. These findings indicate that VuCYP722C or SlCYP722C alone is not adequate for the BC-ring closure with the C-ring stereoselectivity, implying that other factors regulate the stereochemistry of the BC-ring in CYP722C-mediated orobanchol biosynthesis^13, 17^. Despite the progress in identifying the key enzymes for canonical SL formation, no clear actual molecular mechanism of BC-ring closure reaction affording the tricyclic lactone moiety of canonical SL has been proposed.

Elucidating the molecular mechanism of the stereoselective BC-ring closure reaction will clarify the processes through which plants produce these structurally diverse SLs by precisely controlling the C-ring configuration. In this study, we investigated the molecular mechanism of the BC-ring formation leading to orobanchol, using tomato as a model. We reveal the exact function of SlCYP722C and identify a biosynthetic enzyme which catalyzes the stereoselective BC-ring closure reaction occurring after SlCYP722C reaction. We propose a detailed mechanism for the regulation of the C-ring configuration using a computational structural biology approach. These findings facilitate the implementation of artificial manipulation of the SL structures in plants, and provide a basis for the understanding of the role of distinct SLs.

## Results

### Conversion of CLA to 18-oxocarlactonoic acid, the exact function of SlCYP722C

As the first step towards understanding the mechanism of BC-ring closure, we performed a detailed functional analysis of SlCYP722C at the biochemical level. Recombinant SlCYP722C protein was produced in *Escherichia coli*, with truncation of the *N*-terminus transmembrane domain for increased solubility and the *C*-terminus His_6_-tagged for purification (Supplementary Figs. S1, S2A and B). The enzyme reaction mixture of the recombinant SlCYP722C with CLA as a substrate was analyzed using multiple reaction monitoring (MRM) liquid chromatography tandem mass spectroscopy (LC-MS/MS). The recombinant SlCYP722C produced 18-hydroxycarlactonoic acid (18-OH-CLA) (Supplementary Fig. S2C) and a compound with a molecular mass that was 14 Da larger than CLA (CLA+14), in addition to orobanchol diastereomers (Fig. 2A). Considering its molecular weight and involvement in the biosynthesis of orobanchol, CLA+14 was considered a further oxidized derivative of 18-OH-CLA. The methyl esterified CLA+14 by (trimethylsilyl)diazomethane was consistent with the authentic methyl 18-oxocarlactonoate^25^ in LC-MS/MS analysis, identifying CLA+14 as 18-oxocarlactonoic acid (18-oxo-CLA) (Fig. 2B). Moreover, SlCYP722C recognized natural-type (11*R*)-CLA as a substrate, but not (11*S*)-CLA, thereby confirming that the enzyme products have the D-ring in the *R* configuration (Supplementary Fig. S2D and E). The addition of HCl to the enzyme solution at the end of the reaction promoted the formation of orobanchol diastereomers, accompanied by a decrease in 18-oxo-CLA (Fig. 2A). In a separate experiment, the addition of 18-oxo-CLA to a weak acidic buffer (pH 5.8) resulted in its conversion to orobanchol diastereomers over time (Supplementary Fig. S3). These results indicated that 18-oxo-CLA was spontaneously cyclized in a nonenzymatic manner under acidic conditions, supporting our previous study, in which we proposed that the BC-ring closure reaction may be an acid-mediated cascade cyclization based on the conrotatory 4π-electrocyclic reaction (ECR)^25, 26^. Therefore, we conclude that SlCYP722C functions in a two-step oxidation process at the C18 position of CLA to generate 18-oxo-CLA via 18-OH-CLA (Fig. 1D).

**Fig. 2.**
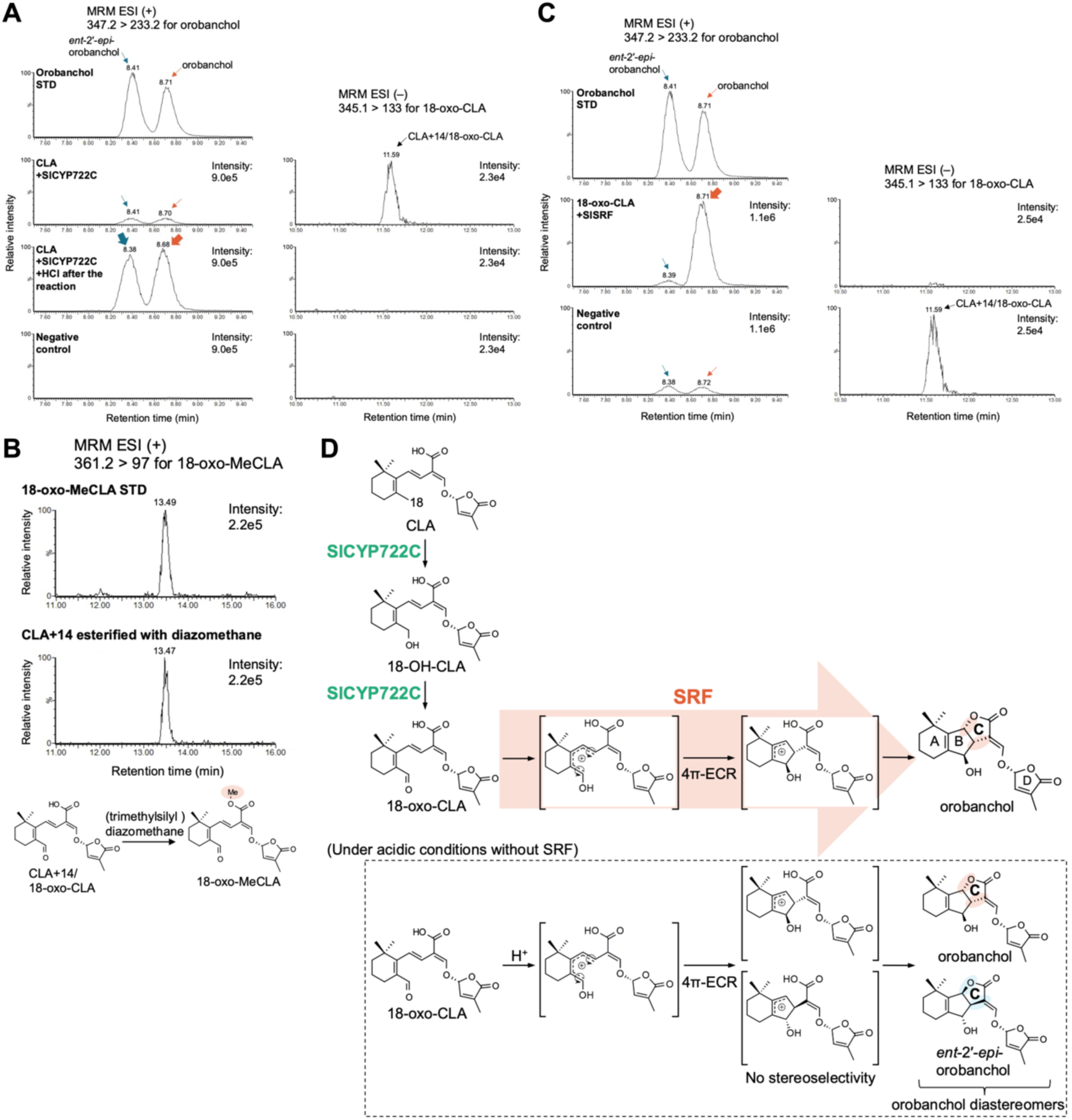
Orobanchol biosynthesis catalyzed by the sequential reaction of SlCYP722C and stereoselective BC-ring-forming factor (SRF). **(A)** *In vitro* enzyme assay of recombinant SlCYP722C; multiple reaction monitoring (MRM) chromatograms of reaction mixtures of recombinant SlCYP722C with carlactonoic acid (CLA) as a substrate. In the negative control, the assay was performed with heat-denatured recombinant SlCYP722C protein. **(B)** Identification of 18-oxocarlactonoic acid (18-oxo-CLA) as a SlCYP722C product, whose molecular mass is 14 Da larger than CLA, by methyl ester derivatization with (trimethylsilyl)diazomethane followed by LC-MS/MS analysis. The retention time and MRM transition of the compound was identical to those of authentic methyl 18-oxocarlactonoate (18-oxo-MeCLA). **(C)** *In vitro* enzyme assay of recombinant SlSRF; MRM chromatograms of reaction mixtures of recombinant SlSRF with 18-oxo-CLA as a substrate. The negative control experiment was performed with heat-denatured recombinant SlSRF protein. **(D)** The putative orobanchol biosynthesis pathway by the sequential reactions of SlCYP722C and SRF downstream of CLA. 18-Oxo-CLA is produced through the reaction by SlCYP722C. The acid-mediated cascade cyclization of 18-oxo-CLA based on 4π-ECR *in flask* generates orobanchol and *ent*-2′-*epi*-orobanchol, whose relative stereochemistry is controlled to be *trans*^25^. In contrast, the conformation of 18-oxo-CLA is controlled within SRF, producing only orobanchol.

### Stereoselective conversion of 18-oxo-CLA to orobanchol by a dirigent domain-containing enzyme *in vitro*

<u>S</u>tereoselective BC-ring-forming factor (SRF), which regulates the stereoselective cyclization of 18-oxo-CLA to generate orobanchol, was inferred to function downstream of SlCYP722C. In tomato and cowpea, as in many other plants, phosphate deficiency increases the expression of SL biosynthesis genes^13, 27, 28^. The gene expression of *SRF* is also expected to upregulate in response to phosphate deficiency. Previously, Wang et al. reported the time-course transcriptional changes in phosphate-deficient and phosphate-replenished tomato roots^29^. The SRF investigation in this study focused on 48 differentially expressed genes at the core of the phosphate response. These genes were upregulated at all timepoints during phosphate deficiency and downregulated with phosphate replenishment^29^ (Supplementary Fig. S4). From this list, we selected the *Solyc01g059900* gene, which encodes for the only dirigent domain-containing protein (DIR; Pfam PF03018) among the 48 genes. Furthermore, we explored the SRF candidates in the gene co-expression network analysis through weighted gene co-expression network analysis (WGCNA)^30^ using our previous cowpea transcriptome dataset (DDBJ Sequence Read Archive, accession no. DRA008222) that led to the discovery of *VuCYP722C*^13^. Notably, this analysis revealed that *Vigun03g413800* and *Vigun03g413900*, homologs of *Solyc01g059900*, along with all known orobanchol biosynthesis genes were included in the same co-expression gene module as the hub genes associated with SL biosynthesis (Supplementary Fig. S5 and Supplementary Data S1).

The DIRs mediate the enantioselective bimolecular coupling, which leads to the stereoselective coupling of coniferyl alcohol radicals in the formation of (+)- or (−)-pinoresinol in lignan biosynthesis^31, 32^. In another case, pterocarpan synthase with a dirigent domain enantiospecifically catalyzed the intramolecular cyclization of 2′-hydroxyisoflavanol to form pterocarpan^33, 34^. The DIR family is classified into six subfamilies (DIR-a, b/d, c, e, f, and g) based on their phylogenetic analyses^35^. Pinoresinol-forming DIRs and pterocarpan synthase belong to the DIR-a and DIR-b/d subfamilies, respectively. Conversely, Solyc01g059900, Vigun03g413800, and Vigun03g413900 belong to the DIR-f subfamily, and the biochemical functions of this subfamily remain elusive (Supplementary Fig. S6). The function of enantioselective substrate conversions by DIR members evoked the possible involvement of DIR as an SRF in the stereoselective formation of orobanchol through the intramolecular BC-ring formation of 18-oxo-CLA.

We biochemically characterized Solyc01g059900 (SlSRF). Reportedly, most DIRs possess a putative signal peptide sequence at the *N*-terminus and are processed into the mature active forms following the truncation of this sequence^34, 36^. For *in vitro* biochemical analysis, recombinant SlSRF was expressed in *E. coli* with the putative signal peptide truncated (i.e., replacing the 24 *N*-terminus residues with Met) and the *C*-terminus His_6_-tagged^34^ (Supplementary Figs. S1 and S7A). Incubation of the purified recombinant SlSRF with 18-oxo-CLA, generated through the reaction of the recombinant SlCYP722C with CLA, resulted in the stereoselective formation of orobanchol with the utilization of 18-oxo-CLA (Fig. 2C and Supplementary Fig. S8). This suggests that Solyc01g059900 acts as an SRF, which catalyzes the stereoselective BC-ring closure reaction of 18-oxo-CLA (Fig. 2D). For cowpea, Vigun03g413900 functioned as an SRF, catalyzed the same reaction to form orobanchol (Supplementary Fig. S9).

### Functional verification of SlSRF *in planta*

To confirm the function of SlSRF in orobanchol biosynthesis *in planta*, we generated two biallelic homozygote *SlSRF*-knockout tomato lines by CRISPR/Cas9-mediated genome editing, ie., *slsrf-1* (with 349 bp deletions of the target regions) and *slsrf-2* (with 13 bp deletions and 1 bp insertion of the target regions) (Fig. 3A). In the root exudates of *slsrf-1* and *slsrf-2*, approximately equal amounts of orobanchol and *ent*-2′-*epi*-orobanchol were detected (Fig. 3B), suggesting that the function loss of SlSRF resulted in the spontaneous generation of orobanchol diastereomers from 18-oxo-CLA. Additionally, we investigated the subcellular localization of SlSRF. The signal peptide of SlSRF fused with mCherry at the *C*-terminus (SlSRF-SP:mCherry) transiently expressed in *Nicotiana benthamiana* leaves revealed that mCherry fluorescence overlapped with enhanced green fluorescent protein fluorescence from the endoplasmic reticulum (ER) marker (Fig. 3C). These results suggest that SlSRF colocalizes to the ER with SlCYP722C, an ER-localized cytochrome P450 (Fig. 3D), facilitating rapid substrate transfer between the two; otherwise, the 18-oxo-CLA produced by SlCYP722C can be cyclized to generate orobanchol diastereomers, as observed in the *in vitro* conversion (Fig. 2 and Supplementary Fig. S3) and root exudates of *slsrf* plants (Fig. 3B). We consequently identified SlSRF, an enzyme belonging to the DIR-f subfamily, catalyzing the stereoselective BC-ring formation in orobanchol biosynthesis.

**Fig. 3.**
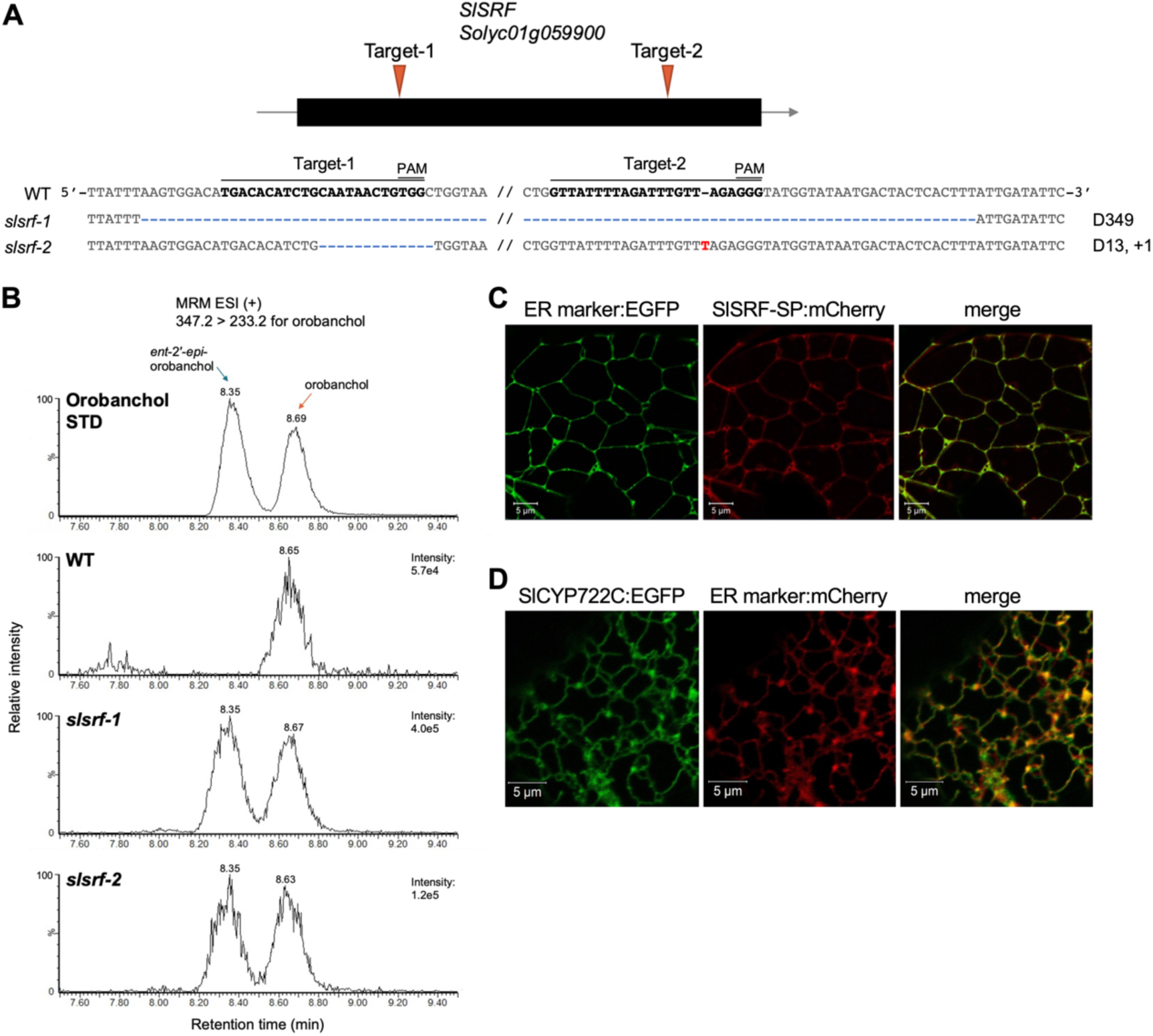
Characterization of SlSRF *in planta*. **(A)** Generation of the *SlSRF*-knockout tomato lines (*slsrf-1* and *slsrf-2*) by CRISPR/Cas9 system. The structure of *SlSRF* with the CRISPR/Cas9 target sites (gRNA Target-1 and Target-2) and sequences are shown. Each target region is shown in bold letters followed by the protospacer adjacent motif (PAM). The number of deleted (D) and inserted (+) nucleotides is indicated on the right side of the sequences. **(B)** Analysis of the orobanchol diastereomers, orobanchol and *ent*-2′-*epi*-orobanchol, in the root exudates of wild-type (WT) and *slsrf*-lines grown under phosphate-deficient conditions. **(C and D)** Subcellular localizations of signal peptide of SlSRF (SlSRF-SP) and SlCYP722C. SlSRF-SP fused with mCherry **(C)** or SlCYP722C fused with enhanced green fluorescent protein (EGFP) **(D)** along with endoplasmic reticulum (ER) markers are transiently expressed in the leaves of *Nicotiana benthamiana.* The results show that SlSRF-SP and SlCYP722 colocalize with the ER markers. Scale bars = 5 μm.

### Computational analysis of the stereoselective ring formation mechanism using a predicted structure of SlSRF

In the molecular mechanism of DIR, a general stereoselective mechanism involving the generation and stabilization of quinone methides (QMs) has been proposed for pinoresinol-forming DIRs and pterocarpan synthase belonging to the DIR-a and DIR-b/d subfamilies, respectively^33, 37^. The biochemical function and molecular mechanism of the DIR-f subfamily were unknown prior to this study. Using a synthetic organic chemistry approach, we proposed an inventive hypothesis where the BC-ring formed through an acid-mediated sequential cyclization process via a conrotatory 4π-ECR mechanism. We further postulated that the BC-ring formation *in planta* proceeds in a similar manner to this acid-mediated cascade cyclization reaction as it occurs “*in flask*”^25, 26^.

We investigated the stereoselective mechanism of SlSRF using a model structure predicted by AlphaFold2^38^ and molecular dynamics (MD) simulations. Monomeric structure of SlSRF was generated with very high pLDDT scores (Fig. 4A), suggesting that the functional domain is located within residues 26–174. Considering the predicted structure and the widely conserved residues among DIRs, we predicted the catalytic residues as D37, D70, Y90, D124, and R131 (Fig. 4B and C). The p*K*a values of D37 and D70 estimated by PROPKA3^39^ were 6.71 and 8.74, respectively. These residues were protonated during the subsequent MD simulations. We docked 18-oxo-CLA into the predicted catalytic site and performed a 30-ns MD simulation with distance restraints (Supplementary Table S1) to refine the complex model. During an additional 100-ns unrestrained MD simulation, the carboxy group of 18-oxo-CLA docked in SlSRF stably formed a salt bridge with R131, and the 18-aldehyde group was oriented to D37 and D70 (Fig. 5A and Supplementary File S1). The latter residues, which were considered proton donors, are conserved in relative enzymes^33^. 18-Oxo-CLA was stably located in the catalytic site. Its major dominant conformation had an approximate C18–C5–C6–C8 dihedral angle (denoted as *φ*) of −30°, with a C8–C18 distance (denoted as *d*) of <3.5 Å (Fig. 5B, C, and Supplementary Movie S1). In particular, the ratio of *φ* under *d* < 3.5, suitable for initiating the subsequent cyclization reaction toward orobanchol (Supplementary Fig. S10), was 99.3%. Meanwhile, the *φ* observed in a unique independent 100-ns MD simulation of 18-oxo-CLA in water displayed a bimodal distribution, with peaks of approximately −30° (43.4%) and 30° (56.6%) (Fig. 5D and Supplementary Movie S2). The latter included the reactant conformation of *ent*-2′-*epi*-orobanchol (Supplementary Fig. S10B). These observations indicate that SlSRF can bind to 18-oxo-CLA tightly and control its conformation.

**Fig. 4.**
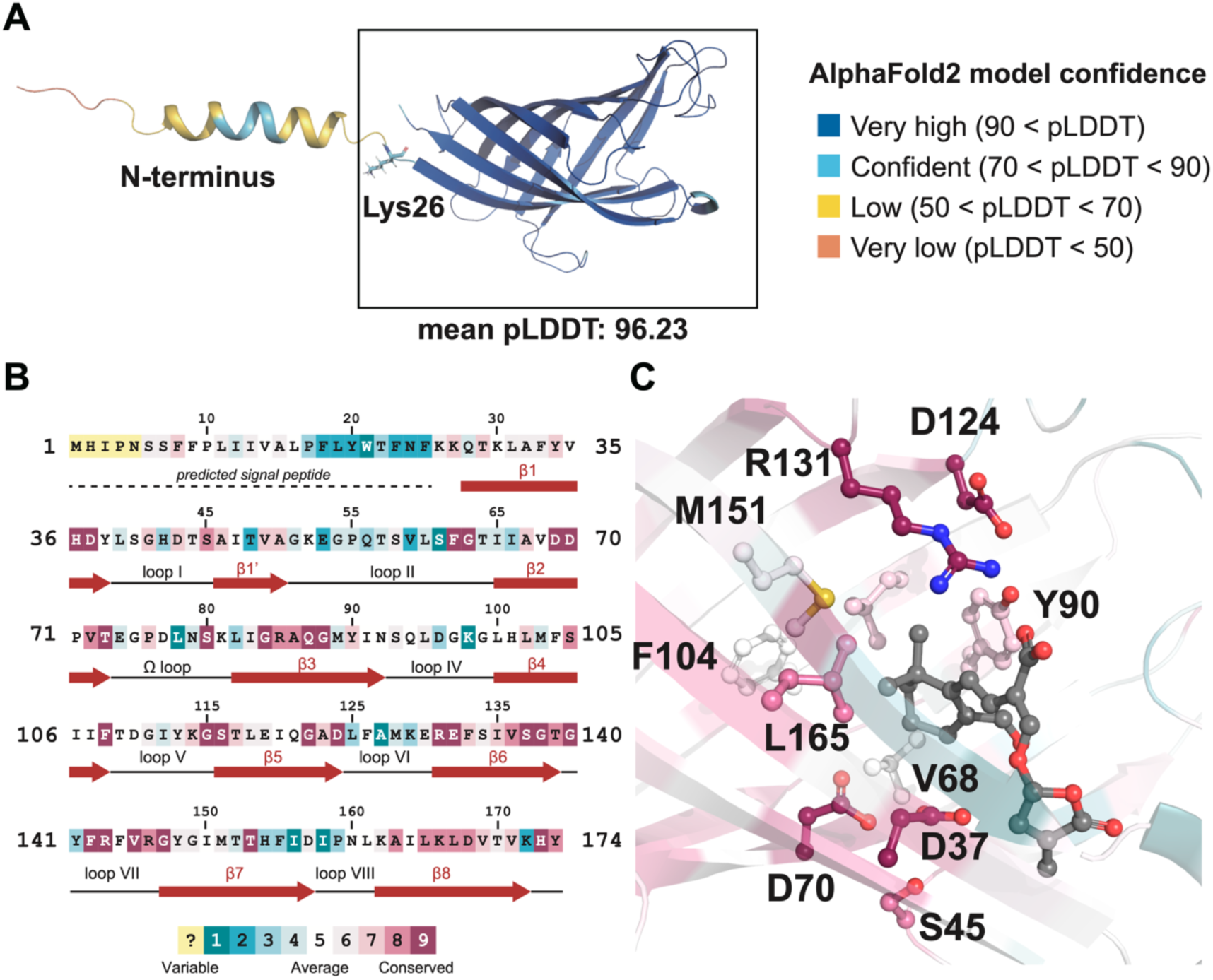
Model structure of SlSRF as predicted by AlphaFold2. **(A)** Predicted monomeric SlSRF structure. The structure is colored by the per-residue pLDDT confidence of AlphaFold2. (**B and C**) Conserved amino acids of SlSRF. The amino acid sequence and predicted structure of SlSRF colored by the conservation scores are shown in (**B**) and (**C**), respectively. The scores were calculated by the ConSurf webserver^46–48^ on Apr. 24, 2023. The 18-oxo-CLA is shown in black. The dark red arrow represents the secondary structure based on its tertiary structure model as predicted by AlphaFold v2.1.0.

**Fig. 5.**
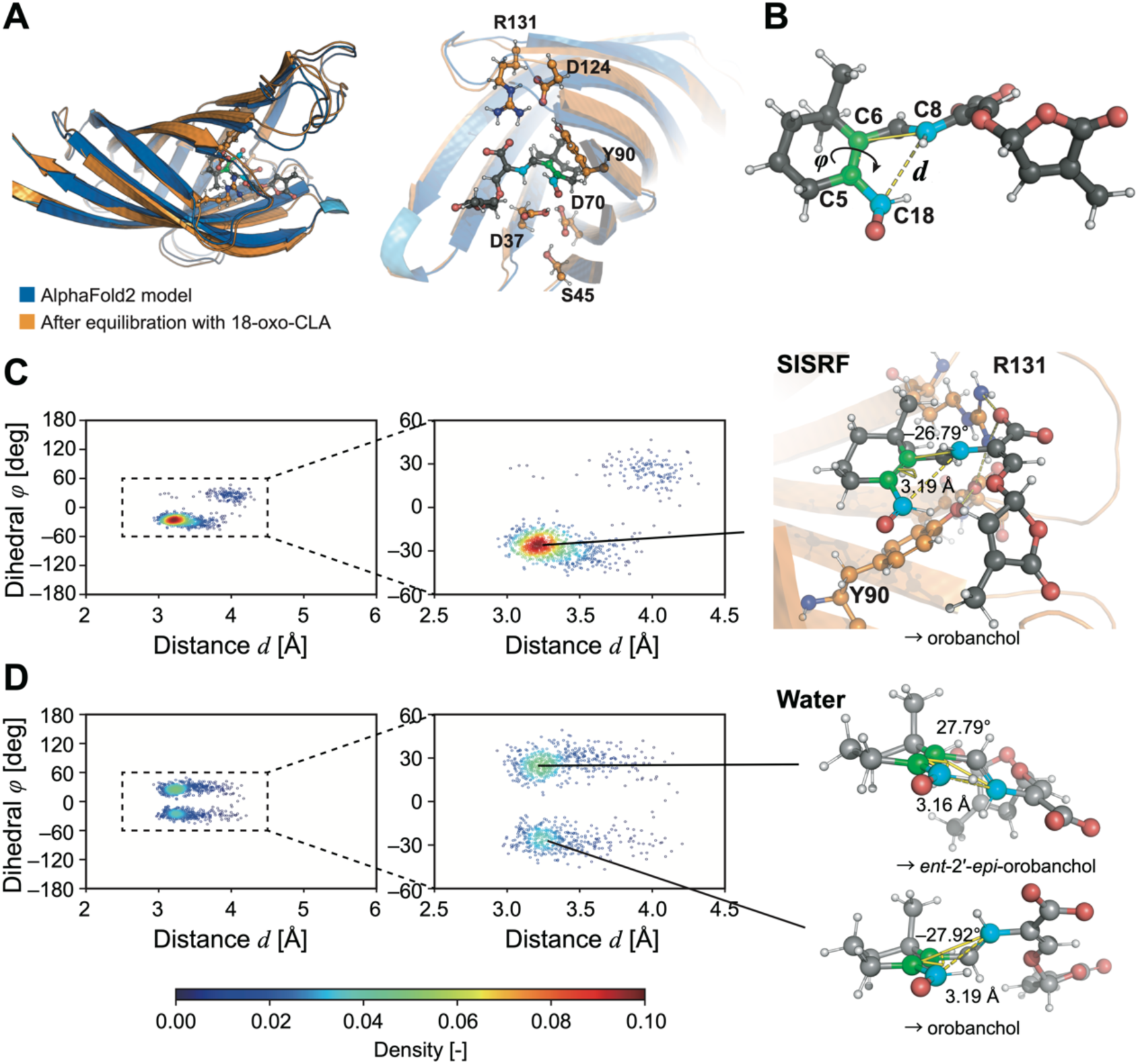
MD simulation analysis based on a model predicted by AlphaFold2. **(A)** Superposition of the AlphaFold2 model (blue) with an equilibrated model in complex with 18-oxo-CLA using MD simulations (orange). A close-up view of the binding pocket is shown in the right panel. **(B)** Definitions of C8–C18 distance (*d*) and C18–C5–C6–C8 dihedral angle (*φ*) in 18-oxo-CLA. C8 and C18 atoms are shown in cyan. C5 and C6 atoms are shown in green. **(C and D)** Differences in conformational distribution of 18-oxo-CLA in complex with SlSRF **(C)** and in water **(D)** observed in each 100-ns MD simulation. The horizontal and vertical axes represent the distance (*d*) and the dihedral angle (*φ*), respectively. The scatter plots are colored according to the density obtained through a Gaussian kernel density estimation.

To gain insight into the catalytic mechanism of SlSRF involved in the stereoselective BC-ring closure reaction, we employed density functional theory (DFT) in addition to our *N*-layered integrated molecular orbital and molecular mechanics (ONIOM) calculations^40–43^ using a snapshot taken from the MD trajectory. As previously stated^25^, the DFT calculations for 18-oxo-CLA suggest that protonation on the 18-aldehyde group is essential to initiate cyclization (Supplementary Fig. S11). In the ONIOM optimization for the reactant state, a proton initially bonded to D37 was spontaneously transferred to the 18-aldehyde group of 18-oxo-CLA through a water molecule (Fig. 6A). Subsequently, the distance between the C8 and C18 atoms was shortened from 2.75 Å to 1.68 Å to yield an intermediate through the transition state, TS1 (Fig. 6A to C). The activation energy (*ΔE*^‡^) of TS1 from the reactant state was 11.5 kcal mol^−1^. The carboxy group of 18-oxo-CLA was then bonded to the C7 atom to yield the product state through another transition state, TS2 (Fig. 6D and E). The activation energy of TS2 from the intermediate state was relatively small (3.9 kcal mol^−1^) but required the removal of the salt bridge between the carboxy group and R131, which was not observed in the reaction pathway *in vacuo* (Supplementary Fig. S10). The calculated reaction energy (*ΔE*) of the product state was −10.9 kcal mol^−1^, and the activation energy from the reactant state was 14.6 kcal mol^−1^. These calculations demonstrated that orobanchol is produced in a conrotatory 4π-ECR-like manner^25, 26^ by SlSRF (Fig. 2D).

**Fig. 6.**
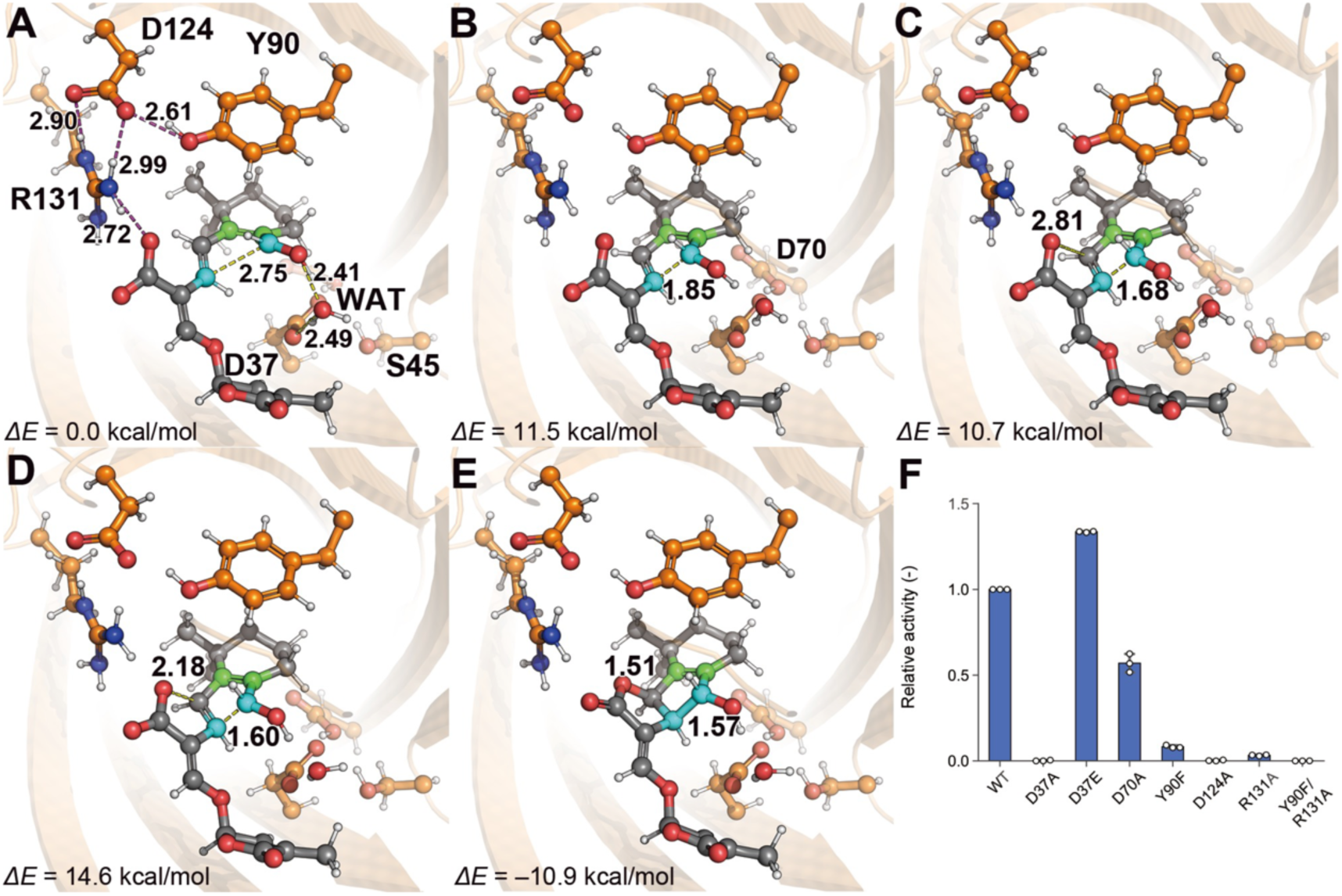
QM/MM analysis of the reaction mechanisms of the BC-ring closure reaction catalyzed by SlSRF. (A–E) Structures of the reactant **(A)**, TS1 **(B)**, intermediate **(C)**, TS2 **(D)**, and product states **(E)** optimized for the SlSRF in complex with protonated 18-oxo-CLA using the ONIOM method. The H-bond network formed by Y90, D124, and R131 are shown in purple dashed lines in the panel **(A)**. Relative energies from the reactant state (*ΔE*) are shown in kcal/mol. **(F)** Relative activity of the SlSRF mutants. Error bars represent standard deviation from the mean (n = 3 replicates).

### Experimental validation of the predicted catalytic residues

Based on the computational results, we performed experimental site-directed mutagenesis to validate the importance of the catalytic residues of SlSRF. Six of seven mutants showed decreased activities, producing less orobanchol than the wild-type (WT) (Fig. 6F), and none of them showed spontaneous cyclization of 18-oxo-CLA to the diastereomer, *ent*-2′-*epi*-orobanchol (Supplementary Fig. S12). The decreases in the enzymatic activities for the D70A, Y90F, D124A, and R131A mutants were in good agreement with the complex model because they impaired hydrogen bond formation to fix 18-oxo-CLA and the cyclization activity (Fig. 6F). Notably, the D124A and Y90F/R131A mutations severely decreased the activities to <1%, indicating that the H-bond network formed by the three residues to fix the carboxy group of 18-oxo-CLA is significant for the enzymatic activity (Fig. 6A). Moreover, the complete loss of activity of the D37A mutant demonstrated that the residue plays a critical role in the initiation of the reaction as a proton donor. Meanwhile, the D37E mutant exhibited enhanced activity over the WT, possibly because the longer side chain made a proton accessible to the 18-aldehyde of the 18-oxo-CLA without intrusion of a water molecule (Supplementary Fig. S13). Taken together, we revealed the detailed cyclization mechanism for the formation of orobanchol from 18-oxo-CLA in SlSRF by experimentally confirming this computationally predicted mechanism.

## Discussion

The mechanism of BC-ring formation in SL biosynthesis is essential for the construction of the basic skeleton of the canonical SL; however, limited information was available related to this process. This work completed the orobanchol biosynthesis pathway by identifying SRF, which belongs to the DIR-f subfamily, as the enzyme which catalyzes the stereoselective cyclization of 18-oxo-CLA to orobanchol (Fig. 2D). Our research, which utilized a combination of experimental and computational methodologies, revealed a detailed catalytic mechanism of stereoselective BC-ring formation for canonical SL by SlSRF in tomato.

All DIRs with previously identified biochemical functions in plants use phenols as their substrates to produce plant specialized phenolic metabolites with QMs as the reaction intermediates. The SlSRF implicated in the biosynthesis of orobanchol exhibits divergent characteristics compared to previously proposed DIRs, as it uses 18-oxo-CLA as a substrate rather than phenolic compounds. However, the conserved amino acid residues within the DIRs, which are also found in SlSRF, suggest the presence of a common functional characteristic responsible for the reaction as a DIR. Notably, the substrate protonation by the aspartate residue, specifically D37 in SlSRF, which is a widely conserved residue among DIRs (Fig. 4B), is a crucial requirement for DIR-mediated catalysis. In our proposed catalytic mechanism, D37 is a donor to protonate the 18-aldehyde group of 18-oxo-CLA through a water molecule, which initiates the sequential cyclization of the B- and C-ring in a 4π-ECR-like manner (Fig. 6). D50 in pterocarpan synthase GePTS1 of *Glycyrrhiza echinata*, a homologous residue of D37 in SlSRF, potentially functions as a proton donor for the hydroxy group of the substrate which leaves as water and eventually forms the QM intermediate^33^. In the pinoresinol-forming DIR AtDIR6 of Arabidopsis, a proposed mechanism involves the 8-8′ coupling of two coniferyl alcohol QM radicals to form a bis-QM intermediate. In this mechanism, a residue that is homologous to D37 in SlSRF (D49 in AtDIR6) serves as a proton donor, protonating the carbonyl oxygen at one end of the bis-QM, which enhances the electrophilicity of the neighboring methide carbon and promotes the cyclization at that half of the bis-QM^37^. Thus, these conserved aspartate residues play a crucial role as proton donors in facilitating both the intra- and intermolecular cyclization reactions. Moreover, Y90, D124, and R131 in SlSRF are suggested to be important for enzyme activity by forming an H-bond network to fix the carboxy group of 18-oxo-CLA. In the GePTS1 mechanism, the role of the conserved R131 (R145 in GePTS1) may also be important for the QM stabilization of the partial negative charge at the carbonyl oxygen^33^, suggesting a similar role for the conserved residues. The function of DIR-f subfamily proteins remains insufficiently explored, and the discovery of the role of the DIR-f protein as SRF in this research represents a significant advance in the understanding of the relevance of the DIR family in plants. Although the biochemical function of most DIRs remains unknown, their participation in the stereoselective biosynthesis of other plant specialized metabolites through a common DIR biochemical mechanism is predictable.

Certain cytochrome P450s, including OsCYP711A2 in rice and the CYP722C subfamily in 5DS-producing plants, are involved in the canonical SL biosynthesis through catalyzing the stereoselective formation of the BC-ring. The ring closure reactions catalyzed by these cytochrome P450s possibly also occur through 4π-ECR^26^ via a molecular mechanism which controls the conformation of the substrate as in SlSRF. Notably, the CYP722C subfamily, which is a crucial enzyme involved in the production of canonical SLs in dicot plants, displays distinct activity depending on the plant in which the different types of canonical SLs are produced. In plant species producing strigol-type SLs, such as cotton, the CYP722C subfamily enzymes catalyze the stereoselective conversion of CLA to 5DS^21–23^. In contrast, we found that SRF is required for the generation of the BC-ring in the subsequent reaction catalyzed by CYP722C subfamily enzymes that convert CLA to 18-oxo-CLA in tomato and possibly other orobanchol-producing dicot plants including cowpea. The loss of SRF function in tomato led to the formation of orobanchol and *ent*-2′-*epi*-orobanchol (Fig. 3A and B), suggesting that SRF exhibits stereoselectivity toward the production of orobanchol excluding the formation of *ent*-2′-*epi*-orobanchol. Although it is unclear why plants such as tomato and cowpea avoid the formation of *ent*-2′-*epi*-orobanchol, the elucidation of the functionality of DIRs belonging to the SRF-clade across various plant species and a comprehensive analysis of *SRF* gene knockout plants will improve the understanding of the significance of the C-ring stereochemistry in canonical SLs.

Our identification of the controlling mechanism of the C-ring configuration, which is the principal determinant for SL diversification, provides a molecular basis for the generation of plants with a tailored SL composition. Structurally different SLs affect the relative abundance of microbial taxa^18, 19, 44^. The germination-stimulating activity toward root parasitic weeds depends on the type of SL, and for some root parasitic weeds, a mixture of SLs with different C-ring configurations significantly decreases their germination rate^17, 45^. Tailored manipulation of SLs would greatly contribute to the understanding of the relationship between distinct SLs and rhizosphere biology, as well as enable the generation of crops that recruit beneficial microbes and diminish the incidence of root parasitic weeds by suppressing their germination. Finally, we demonstrate usefulness of these complementally approaches between biochemistry and computational structural biology addressing the molecular mechanism of the stereoselective formation of SL, and its applicability to other plant biosynthetic systems for which enzyme structures have not been experimentally determined.

## Methods

### Chemicals

Authentic samples of CLA, 18-oxo-MeCLA, and orobanchol diastereomers were prepared as previously described^13, 25, 49, 50^.

### Analysis of SLs

The SLs were analyzed using LC-MS/MS system (Waters) comprising Acquity UPLC H-Class and an Acquity TQD tandem mass spectrometer under analytical conditions as previously described^13^. Chromatographic separation was performed with an ODS column (COSMOSIL 2.5C18-MS-II, 100 × 2.0 mm i.d., 2.5 µm; Nacalai Tesque, Kyoto, Japan) at a column oven temperature of 30°C. The elution was performed in a linear gradient system of MeOH–H_2_O with 0.1% formic acid (50:50–100:0 in 20 min) at a flow rate of 0.2 mL min^−1^. Multiple reaction monitoring (MRM) was used to detect SLs, and MRM transitions are summarized in Supplementally Table S2. Data acquisition and analysis were performed using MassLynx 4.1 software (Waters).

For analysis of SLs in root exudates of tomato, tomato plants were grown hydroponically as follows. Seeds of tomato were sown on Murashige-Skoog agar plates and grown at 23°C and a 16-hour light/8-hour dark photoperiod for 7 days. The seedlings were transferred to conical tubes and grown hydroponically in 50-mL of half-strength Hoagland nutrient solution without phosphate under the same conditions as above. After 3 days, the culture medium was replaced with fresh one and plants were grown for a further 7 days. Each filtrate of the hydroponic solution was subjected to purification with Oasis HLB Vac Cartridge (Waters) as previously described^51^.

### Production of recombinant SlCYP722C by heterologous expression in *E. coli* and purification

To produce SlCYP722C in a soluble form, the *N*-terminus 34 residues containing the transmembrane α-helix was replaced with two residues, Met-Ala, as reported previously^52^. The transmembrane region was predicted by the program TMHMM (Supplementary Fig. S1)^53^. The SlCYP722C sequence with the *N*-terminus truncation and the *C*-terminus His_6_-tag immediately followed by a stop codon (ΔN34-SlCYP722C-His_6_) was codon-optimized for expression in *E. coli* and synthesized by Eurofins Genomics (Supplementary Table S3). The DNA fragment was inserted into the cloning site of pColdIII expression vector (TaKaRa Bio, Shiga, Japan). The constructed vector and the chaperone co-expression plasmid pGro12^54^ were introduced into *E. coli* strain BL21 (DE3). The transformed *E. coli* was grown at 37°C in Terrific Broth medium supplemented with 100 μg mL^−1^ carbenicillin and 50 μg mL^−1^ kanamycin.

Overnight cultures were diluted into Terrific Broth media including 100 μg mL^−1^ carbenicillin and 50 μg mL^−1^ kanamycin, and 0.5 mg mL^−1^ L-arabinose (for induction of GroES/GroEL expression), and continued to incubate until the OD600 reached 0.7. After the cultures were iced for 30 min, 0.1 mM Fe(III) citrate and 1 mM 5-aminolevulinic as a haem biosynthesis precursor were added; gene expression was induced by the addition of 0.1 mM isopropyl β-D-thiogalactoside and cells were incubated another 48 h at 15°C. Collected cells were washed with phosphate-buffered saline and resuspended in the lysis buffer (50 mM potassium phosphate buffer (pH 7.4), 500 mM KCl, 5 mM MgCl_2_, 0.1 mM EDTA, 0.1 mM DTT, 20 % (v/v) glycerol, 1.0% (v/v) tween20, 1.0% (w/v) sodium cholate, 5 mM ATP, and 25 mM imidazole) and disrupted by sonication. After centrifugation at 10,000 *g* for 30 min at 4°C, the supernatant was applied on Ni Sepharose column (Cytiva) equilibrated with the lysis buffer. Bound proteins were washed twice with 10 column volumes of the lysis buffer and eluted with 5 column volumes of the elution buffer (50 mM potassium phosphate buffer (pH 7.4), 500 mM KCl, 0.1 mM EDTA, 0.1 mM DTT, 20% (v/v) glycerol, 1.0% (v/v) tween20, 1.0% (w/v) sodium cholate, and 500 mM imidazole). The eluent was diluted with buffer A (50 mM potassium phosphate buffer (pH 7.4), 0.1 mM EDTA, 0.1 mM DTT, and 20 % (v/v) glycerol), and concentrated using a 10 kDa cut-off Amicon Ultra concentrator (Merck). Concentrated and desalted protein solution was applied on a HiTrap DEAE column (Cytiva) equilibrated with buffer B (50 mM potassium phosphate buffer (pH 7.4), 0.1 mM EDTA, 0.1 mM DTT, 20% (v/v) glycerol, and 0.5% (v/v) tween20). The column was washed with 5 column volumes of buffer B. Pass-through and wash fractions were collected and then applied on CM-Sepharose column (Cytiva) equilibrated with buffer B. The column was washed with 10 column volumes of buffer C (50 mM potassium phosphate buffer (pH 7.4), 0.1 mM EDTA, 0.1 mM DTT, 20% (v/v) glycerol, and 1.0% (w/v) CHAPS) and subsequently 10 column volumes of buffer C with 50 mM KCl. Proteins were eluted with 5 column volumes of buffer C with 400 mM KCl. Finally, elution fraction was collected and concentrated using a 10 kDa cut-off Amicon Ultra concentrator (Supplementary Fig. S2A). The active recombinant SlCYP722C was quantified based on the carbon monoxide (CO)-difference spectra using an extinction coefficient (Σ = 91 mM^−1^ cm^−1^) (Supplementary Fig. S2B)^55^.

### Biochemical characterization of SlCYP722C

For *in vitro* enzyme activity assay, the reaction mixture consisted of 50 mM potassium phosphate (pH 7.4), 80 nM purified recombinant SlCYP722C, 1 unit/mL Arabidopsis NADPH–CYP reductase^56^, 2.5 mM NADPH, and 10 μM CLA as a substrate in a total volume of 100 μL. Reactions were initiated by the addition of NADPH and performed at 30°C for 60 min. The reaction products were extracted twice using an equivalent volume of ethyl acetate; for HCl treatment, 10 μl of 5N HCl was added after the reaction. The collected organic phase was evaporated. The residues were dissolved in acetonitrile and analyzed by LC-MS/MS.

To obtain 18-OH-CLA by sorghum (*S. bicolor*) SbCYP711A31^57^, the *in vitro* enzyme assay was conducted as follows. Recombinant SbCYP711A31 was obtained in a baculovirus-insect cell expression system as previously described^13^. Briefly, the SbCYP711A31 cDNA was ligated into the *Bam*HI/*Sal*I site of the pFASTBac1 vector (Invitrogen). Recombinant bacmid DNA was then produced by transforming *E. coli* DH10Bac competent cells (Invitrogen). Preparation of recombinant bacmid DNA and transfection of *Spodoptera frugiperda* 9 (*Sf*9) cells were performed according to the manufacturer’s instructions (Invitrogen). SbCYP711A31 was expressed in *Sf*9 cells and microsomal fractions of the insect cells expressing SbCYP711A31 were obtained as described previously^58^. The microsomal fraction was mixed with the reaction solution of the composition shown above instead of the recombinant SlCYP722C, and incubated at 30°C for 60 min.

### Methyl ester derivatization of CLA+14 produced by SlCYP722C

The SlCYP722C enzyme reaction product containing CLA+14 was dissolved with 500 μL of methanol/dichloromethane (20:80, v/v), and 250 μl of (trimethylsilyl)diazomethane was added and allowed to react for 30 minutes at room temperature to esterify free carboxylic acids. The reaction solution was then dried and dissolved in acetonitrile for LC-MS/MS analysis.

### Evaluation of the stability of 18-oxo-CLA

The SlCYP722C enzyme reaction products, extracted with ethyl acetate and dried, were dissolved with a small volume of acetonitrile and dispensed with water. Equal volumes of 100 mM potassium phosphate buffer (pH 7.4 or 5.8) were added to each aliquot and incubated at 25°C. Immediately after the addition of buffer, 2, 4, and 6 hours later, the samples were extracted twice with ethyl acetate containing 200 nM 2′-*epi*-strigol as an internal standard, then dried and dissolved in acetonitrile for LC-MS/MS analysis.

### Expression data preparation and weighted gene co-expression network analysis using cowpea RNA-seq dataset

The previous our raw reads of cowpea RNA-seq dataset (DDBJ Sequence Read Archive, accession no. DRA008222)^13^ were processed by trimming adaptor and low-quality sequences using Fastp software^59^ with default parameters. Using HISAT2^60^, pre-processed fastq files were mapped to the cowpea reference genome^61^. The SAM files obtained were sorted and converted into BAM files using Samtools^62^. The FPKM values were quantified for each sample to measure the gene expression levels using StringTie^63, 64^. Subsequent analysis was performed for 20,480 genes with the FPKM values greater than 0 in all samples.

A weighted gene co-expression network was constructed using WGCNA package in R^30^. Network construction and module detection were implemented by ‘blockwiseModules’ function with a soft threshold power of 22 which was determined using ‘pickSoftThreshold’ function, a merge cut height of 0.25, and a minimum module size of 30, resulting in 39 co-expression modules were identified (Supplementary Fig. S5). Individual genes within the modules were ranked by module membership defined as the correlation between the expression profile and the module eigengene (in other words, a measure of how similar the expression profile of a gene is to that of the module). Screening for genes with high module membership has been shown to be an effective strategy for identifying genes of biological interest^65^. The tan co-expression module containing 388 genes included all known genes involved in orobanchol biosynthesis in cowpea, namely *VuD27 (Vigun01g137900)*, *VuCCD7 (Vigun05g149500)*, *VuCCD8 (Vigun09g192400)*, and *VuCYP711A (Vigun09g224400)*, and *VuCYP722C (Vigun03g264300)*, with high module membership; thus, the top 100 genes with high module membership in the tan co-expression module were extracted as orobanchol biosynthesis-associated hub genes (Supplementary Dataset S1).

### Phylogenetic analysis

To characterize the phylogenetic relationships of SRFs, the amino acid sequences of DIRs from many plant species previously described^33, 66–68^, including *Arabidopsis thaliana*, *Forsythia intermedia*, *Glycine max*, *Glycyrrhiza echinata*, *Gossypium hirsutum*, *Linum usitatissimum*, *Oryza sativa*, *Picea glauca*, *Pisum sativum*, *Schizandra chinensi*, and *Thuja plicata*, as well as those of paralogs of SlSRF and VuSRF selected by blastp search were analyzed using MUSCLE^69^. A phylogenetic tree was constructed in MEGA X^70^ by the maximum-likelihood method using the substitution model LG+G with 1000 bootstrap replicates and drawn in iTOL^71^.

### Expression and purification of recombinant SRFs

The sequences of *Solyc01g059900* (*SlSRF*) in tomato and *Vigun03g413800* and *Vigun03g413900* (*VuSRF*) in cowpea were PCR-amplified using the primers listed in Supplementary Table S4. Each primer was designed with the *N*-terminus putative signal peptide truncated. The transmembrane region was predicted by the program TMHMM^53^ (Supplementary Fig. S1). For expression as the *C*-terminus His_6_-tag fused protein, the resultant PCR products were inserted into the *Nde*I and *Kpn*I restriction sites of the modified pColdIII expression vector (TaKaRa Bio, Shiga, Japan) containing the His_6_-tag sequence with stop codon (5’-catcaccatcaccatcactag-3’) immediately after the *Kpn*I site of the multicloning site. The constructed vector was introduced into *E. coli* strain BL21 (DE3). The transformed *E. coli* was grown at 37°C in Terrific Broth medium supplemented with 100 μg mL^−1^ carbenicillin.

Overnight cultures were diluted into Terrific Broth media including 100 μg mL^−1^ carbenicillin and continued to incubate until the OD600 reached 0.7. After the cultures were iced for 30 min, gene expression was induced by the addition of 0.1 mM isopropyl β-D-thiogalactoside and cells were incubated another 48 h at 15°C. Collected cells were washed with phosphate-buffered saline.

For purification of the recombinant SlSRF, the SlSRF expressing cells were resuspended in the lysis buffer (50 mM potassium phosphate buffer (pH 7.4), 500 mM KCl, 0.1 mM EDTA, 0.1 mM DTT, 20 % (v/v) glycerol, 1.0 % (w/v) CHAPS, and 50 mM imidazole) and disrupted by sonication. After centrifugation at 10,000 *g* for 30 min at 4°C, the supernatant was applied on TALON spin column (TaKaRa Bio, Shiga, Japan) equilibrated with the lysis buffer. After incubation at room temperature for 5 min, the column was centrifuged at 100 *g* for 1 min, and pass-through was discarded. Bound proteins were washed twice with 5 column volumes of the lysis buffer and eluted twice with 2 column volumes of the elution buffer (50 mM potassium phosphate buffer (pH 7.4), 500 mM KCl, 0.1 mM EDTA, 0.1 mM DTT, 20 % (v/v) glycerol, 1.0 % (w/v) CHAPS, and 500 mM imidazole). The eluent was diluted with buffer A, and concentrated and desalted using a 10 kDa cut-off Amicon Ultra concentrator (Merck) (Supplementary Fig. S7A).

For site-directed mutagenesis, SRF mutants were prepared using PrimeSTAR Mutagenesis Basal Kit according to the manufacturer’s instructions (TaKaRa Bio, Shiga, Japan), and the mutations were confirmed by sequencing. The primers used for the experiments are listed in Supplementary Table S4. The pColdIII vector containing SlSRF described above was used as the template for PCR. Resulting mutated plasmids were transformed into *E. coli* strain BL21 (DE3). The SRF mutants were expressed and purified as described above (Supplementary Fig. S7B).

Protein content was estimated using Pierce BCA Protein Assay Kit according to the manufacturer’s instructions (Thermo Fisher Scientific).

For Vigun03g413800 and Vigun03g413900 expressing cells, the cells were resuspended in buffer A (50 mM potassium phosphate buffer (pH 7.4), 0.1 mM EDTA, 0.1 mM DTT, 20 % (v/v) glycerol), disrupted by sonication, and centrifuged at 10,000 *g* for 30 min at 4 °C and the supernatants were used as crude protein for subsequent assays (Supplementary Fig. S9A).

### *In vitro* enzyme activity assay for SRFs

The substrate 18-oxo-CLA was prepared at a time of use as follows. As described above, the SlCYP722C enzyme reaction products, extracted with ethyl acetate and dried, were dissolved with a small volume of acetonitrile and dispensed with 50 mM potassium phosphate buffer (pH 7.4). The reaction mixture consisted of 50 mM potassium phosphate (pH 7.4) and 40 nM purified recombinant SlSRF, or crude protein of Vigun03g413800 or Vigun03g413900, and 18-oxo-CLA as a substrate in a total volume of 100 μL. Reactions were performed at 30°C for 30 min. The reaction products were extracted twice using an equivalent volume of ethyl acetate containing 200 nM 2′-*epi*-strigol as an internal standard. The collected organic phase was evaporated. The residues were dissolved in acetonitrile and analyzed by LC-MS/MS.

For the site-directed mutagenesis, enzyme reactions were performed as above. The amount of orobanchol produced by the enzymatic reaction was estimated by calculating the relative peak area of orobanchol to that of internal standard after the reaction, and subtracting from this value the relative peak area of orobanchol in the negative control (assay with heat-denatured protein). The relative activity was expressed by comparing the amount of orobanchol produced by the wild-type and each mutant.

### Generation of *SlSRF* knockout tomato plants

The *SlSRF* (*Solyc01g059900*) knockout tomato (*S. lycopersicum* cv. Micro-Tom) plants were generated by targeted genome editing with the CRISPR/Cas9 system, as described previously^13^. *SlSRF* gene was targeted by two guide RNAs (gRNAs) (Fig. 3A). To design the gRNA targets with low off-target effects, the web tool CRISPR-P v2.0^72^ (http://cbi.hzau.edu.cn/cgi-bin/CRISPR2/CRISPR) was used to select two gRNAs (Target-1: 5′-tgacacatctgcaataactgtgg-3′ and Target-2: 5′-gttattttagatttgttagaggg-3′) which were estimated to have significantly low off-target scores. The DNA fragment composed of the gRNA and transfer RNA (tRNA) scaffolds between both target sequences was generated by PCR using pMD-gtRNA containing gRNA and tRNA scaffolds as a template and primer sets containing restriction enzyme *Bsa*I sites (5′-ttgggtctcgtgcagtgacacatctgcaataactggttttagagctagaaatagca-3′ and 5′-ttgggtctccaaactctaacaaatctaaaataactgcaccagccgggaatcgaa-3′). The unit containing two gRNAs-tRNA was then inserted into the *Bsa*I site of multiplex CRISPR-Cas9 vector pMgP237-2A-GFP^73^ using Golden Gate cloning methods to generate the CRIPR-Cas9 vector pMgP237-SlSRFKO. The vector was introduced into *Agrobacterium tumefaciens* strain EHA105 by electroporation. Tomato plants were transformed using *A. tumefaciens* EHA105 cells harboring pMgP237-SlSRFKO as reported previously^13^. Transformants were selected using kanamycin and genomic PCR targeting a partial region of T-DNA region integrated into the genome with the primers 5′-ggcccctgggaatctgaaag-3′ and 5′-ggaagaagaaatcgatctggaaaattttgc-3′.

The CRISPR-mediated mutations were identified by amplifying the DNA region that contains the gRNA target sites with primer sets used for cloning of *SlSRF*, and by sequencing the resulting PCR fragment. The T0 lines carrying the targeted mutations were selected to generate T1 and T2 progeny by self-pollination, resulting in two T2 biallelic homozygote *SlSRF* knockout lines, *slsrf-1* with 349 bp deletions, and *slsrf-2* with13 bp deletions and 1 bp insertion of the target regions, respectively (Fig. 3A).

### Sub-cellular localization analysis

For examining the sub-cellular localization of SlSRF, a cDNA encoding putative signal peptide of SlSRF (SlSRF-SP) (AA 1–24) or a full-length *SlCYP722C* cDNA without stop codon was fused with the *C*-terminus mCherry or enhanced green fluorescent protein (EGFP), and each was cloned into *Nde*I/*Sal*I sites of the pRI201-AN vector (TaKaRa Bio, Shiga, Japan) to express under the control of the 35S promoter. For the endoplasmic reticulum (ER) marker construction, the DNA fragment created by inserting HDEL sequence at the *C*-terminus of EGFP or mCherry genes and adding the signal peptide of AtWAK2 at the *N*-terminus^74^ was inserted into *Nde*I/*Sal*I sites of the pRI201-AN vector. The pRI201-AN vector with *p19* coding sequence inserted at the *Nde*I/*Sal*I sites was used as RNA silencing suppressor. *Agrobacterium tumefaciens* strain GV3101 was transformed with the constructed plasmid by electroporation and cultured separately as described previously^50^.

Agroinfiltration was performed using leaves of *N. benthamiana* grown at 25°C for 4 weeks under long-day conditions (16 h light/8 h dark) as described previously^50^. The SlSRF-SP or SlCYP722C, fused with the *C*-terminus fluorescent protein, was transiently co-expressed with ER marker and p19. After infiltration, the plants were incubated for 2 days under the same conditions. *N. benthamiana* leaf pieces were directly mounted on glass slides with a drop of water and then covered with glass covers. mCherry and EGFP were excited at 555 nm and 488 nm, respectively, and observed in the range of 570–600 and 490–520 nm, respectively, using a LSM700 laser scanning confocal microscope (Carl Zeiss).

### Structure prediction of SlSRF

The protein structure of SlSRF was predicted using AlphaFold v. 2.1.0^38^ with ‘monomer-ptm’ model and all genetic databases used at CASP14. The amino acid sequence is downloaded from accession number A0A3Q7EF24 of UniprotKB database. Hhblits^75^ and HHSearch^76^ from hh-suite v.3.3 were used for generation of multiple sequence alignment (MSA) and for the template search with PDB70 database released on Oct. 27, 2021, respectively. The average per-residue pLDDT score of predicted functional domain (residues 26–174) was 96.23 (60.72 for residues 1–25), indicating that the region was predicted with high confidence (Fig.4A). The region was then truncated and used for subsequent molecular dynamics (MD) simulations with its substrate, 18-oxo-CLA.

### Molecular dynamics (MD) simulations

The truncated SlSRF model (residues 26–174) was used as the initial coordinates for MD simulations. According to the best output structure (“ranked_0”) of AlphaFold2, histidine residues at positions 42, 101, and 154 were predicted to be δ-protonated, with the remainder in the ε-protonated state. The geometry of 18-oxo-CLA was fully optimized at the B3LYP/6-31G+(d,p) level and their partial charges were obtained by restrained electrostatic potential (RESP) using HF/6-31G(d) single-point calculations on the optimized geometry using Gaussian 16 Rev C.01^77^. The ff14SB force field^78^ and the general AMBER force field 2 (GAFF2)^79^ were used for the protein and the substrate, respectively. The ANTECHAMBER module was used to parameterize the substrate^80^.

The substrate was docked into the model structure by hand using PyMOL 2.5.0. The ligand was initially placed with reference to a putative model of GePTS1 in complex with (3*S*, 4*R*)-7,2′-dihydroxy-4′-methoxyisoflavanol depicted in the paper^33^, because HHSearch predicted GePTS1 as the most similar protein in the PDB70 database. Considering widely conserved residues among DIRs, D37, D70, Y90, D124 and R131 are estimated to be the catalytic residues, and the aldehyde group of the substrate should be oriented to D37 and D70 to be protonated. Since the p*K*a values of D37 and D70 estimated by PROPKA3 were 6.71 and 8.74, respectively, these residues are assigned to be protonated during subsequent MD simulations.

The prepared complex structure was fully solvated with the TIP3P water model^81^ and 60 Na^+^ ions in a cubic periodic box with an edge length of 80 Å and neutralized by Cl^−^ counter ions using the AMBER LeaP module. Van der Waals interactions were cut off beyond 10 Å. The particle mesh Ewald (PME) method^82^ was used to calculate electrostatic interactions. The system was first relaxed using 200 steps of steepest descent minimization with a 1,000 kcal mol^−1^ Å^−2^ position restraints applied to the heavy atoms of the protein. Subsequently, the restraints were removed and the entire system was subjected to 200 steps of steepest descent minimization. Next, to gradually heat the system, MD simulations over 1 ns in duration at a temperature of 300 K under the *NPT* ensemble were performed. During the equilibration, the SHAKE algorithm^83^ was used to constrain the bonds, including hydrogen atoms. The integration time step was set to 2 fs. The Berendsen weak coupling algorithm^84^ was used to maintain a constant temperature and pressure. All energy minimization, equilibration, and production runs were performed using the PMEMD module of AMBER 20^85^.

### QM/MM (ONIOM) computational scheme

Snapshots taken from the production run were used as initial coordinates of our own *N*-layered integrated molecular orbital and molecular mechanisms (ONIOM)^40^ models in Gaussian 16 Rev C.01. Water molecules more than 5.0 Å away from the protein–ligand atoms were removed to reduce the calculation cost. Two-layer extrapolated ONIOM method was applied to the complex model. The QM and MM layers were treated using the B3LYP/6-31+G(d,p)^86, 87^ level of theory and AMBER ff14SB force field/GAFF2, respectively. We created a QM model, composed of the side chains of D37, S45, D70, Y90, D124, R131, a water molecule, and the ligand, 18-oxo-CLA. To validate the difference of reactivity, we prepared two states of 18-oxo-CLA; one is protonated at the 18-formyl group and the other is neutral. The covalent boundary between the QM and MM layers was capped by hydrogen link atoms. All intermediates and transition states were optimized using the ONIOM mechanical embedding (ME) scheme.

In the first structural optimization, all atoms in the model were not fixed to reduce steric repulsion. Partial charges in the QM layer were then updated to reduce the error caused by the use of fixed charges, as performed by Tao *et al*^88^.The structural optimization was then performed again with residues within 7 Å of the QM layer allowed to move with the rest fixed. The TS coordinates was tested through intrinsic reaction coordinate (IRC) calculations to confirm that they connected the correct reactant and product structures. The electronic embedding (EE) scheme^43^ was used for single-point energy calculations at the ONIOM(B3LYP/6-31++G(d,p):AMBER) level of theory on the ONIOM-ME optimized geometries.

## Data availability

The data supporting the findings of this study are available within the article and its Supplementary Information and are publicly accessible via cited repositories or are available from the corresponding author on reasonable request.

## Acknowledgments

We are grateful to Prof. Osakabe at Tokushima University for providing a genome editing vector, pMgP237-2A-GFP. We would like to thank Enago (www.enago.jp) for the manuscript review and editing support. This work was supported by Science and Technology Research Partnership for Sustainable Development (SATREPS), Japan Science and Technology Agency (JST)/ Japan International Cooperation Agency (JICA) (No. JPMJSA1607 to Y.S.), by ACT-X, JST (No. JPMJAX20BM to T.W.), by PRESTO, JST (No. JPMJPR22DA to T.W.), by Japan Society for the Promotion of Science (JSPS) KAKENHI (Nos. 25292065 to Y.S. and 20K15459 to T.W.), by Research Support Project for Life Science and Drug Discovery (Basis for Supporting Innovative Drug Discovery and Life Science Research (BINDS)) from AMED under Grant Number JP22ama121027 (T.T., support number 4143).

## Contributions

Conceptualization: T.W., Y.M., H.T., and Y.S. Methodology: M.H., T.W., Y.M., and D.O. Investigation: M.H., T.W., Y.M., N.S. T.S., and K.I. Visualization: M.H., T.W., and Y.M. Funding Acquisition: T.W., T.T., K.S., and Y.S. Project administration: T.T. and Y.S. Writing – original draft: M.H., T.W., and Y.M. Writing–review and editing: M.H., T.W., Y.M., A.O. T.T., K.S., M.M., H.T., and Y.S.

## Supplementary Materials

**Supplementary Fig. S1. *In silico* analysis of the SlCYP722C and SRF peptides.** The plot shows the posterior probabilities of inside, outside, and transmembrane helix for each amino acid residue.

**Supplementary Fig. S2. Heterologous expression of the *N*-terminus truncated SlCYP722C and its activity. (A)** SDS-PAGE of the recombinant SlCYP722C protein. The protein was expressed as the *N*-terminus truncated and the *C*-terminus His_6_-tagged form in *E. coli* and purified using a series of affinity columns. M, protein marker; a, crude protein; b, purification after His_6_-tag affinity chromatography; c, purification after DEAE Sepharose anion exchange chromatography; d, final purified protein after CM Sepharose cation exchange chromatography. **(B)** Carbon monoxide (CO)-difference spectra of the recombinant SlCYP722C protein. **(C)** Identification of 18-hydroxy-CLA (18-OH-CLA) as the SlCYP722C product. The reaction product of SlCYP722C with 16 Da larger molecular mass than CLA was identified as 18-OH-CLA by comparing the reaction product of SlCYP722C with that of sorghum (*Sorghum bicolor*) SbCYP711A31, which is known to catalyze the conversion of CLA to 18-OH-CLA^57^. **(D)** Structures of (11*R*)-CLA and (11*S*)-CLA. **(E)** Substrate specificity of SlCYP722C. In the case of (11*R*)-CLA as a substrate, 18-oxo-CLA was formed with the decrease of substrate, whereas the reaction did not proceed with (11*S*)-CLA.

**Supplementary Fig. S3. Formation of orobanchol diastereomers by spontaneous cyclization of 18-oxo-CLA in buffer solution.** The relative amounts of 18-oxo-CLA and orobanchol diastereomers to the internal standard over time differ in sodium potassium buffer at different pH. The decrease in 18-oxo-CLA and the increase in orobanchol diastereomers are more remarkable under acidic conditions.

**Supplementary Fig. S4. 48 Differentially expressed genes at the core of the phosphate response in tomato.** Figure adapted and partially modified from Wang et al^29^ under the terms of a Creative Commons Attribution 4.0 International License (CC BY 4.0). Venn diagram of up-regulated differentially expressed genes (DEGs) at different time points of phosphate deficiency and down-regulated DEGs by phosphate replenishment.

**Supplementary Fig. S5. WGCNA co-expression network and module-trait correlation analysis. (A)** Hydroponic culture conditions of cowpea. Figure adapted and partially modified from our previous paper^13^ under the terms of a Creative Commons Attribution NonCommercial Licenses 4.0 (CC BY-NC 4.0). **(B)** Dendrogram plot with color annotation. **(C)** Correlations of WGCNA modules with hydroponic culture conditions (traits) and corresponding P-values. Each row corresponds to each of the 39 co-expression modules and the columns correspond to six traits. The color of each cell indicates the correlation coefficient between the module and traits; the color scale on the right shows module-trait correlation from –1 (blue) to 1 (red).

**Supplementary Fig. S6. Phylogenetic tree of DIR sequences.** The amino acid sequences of tomato SlSRF (Solyc01g059900.4.1), cowpea VuSRF (Vigun03g413900), and their paralogs and DIRs from other species were used to conduct the phylogenetic analysis in MEGA X using the maximum likelihood method. DIRs were classified into six groups, DIR-a to DIR-f, based on their phylogenetic relationships with reference to previous report^33^ and are indicated by different colors in the phylogenetic tree. The clade containing the SRFs is shaded in pink. At, *Arabidopsis thaliana*; Fi, *Forsythia intermedia*; Ge, *Glycyrrhiza echinata*; Gh, *Gossypium hirsutum*; Gm, *Glycine max*; Lu, *Linum usitatissimum*; Os, *Oryza sativa*; P, *Picea glauca*; Ps, *Pisum sativum*; Sc, *Schizandra chinensis*; Tp, *Thuja plicata*.

**Supplementary Fig. S7. Purification of recombinant SlSRF protein. (A)** SDS-PAGE of the recombinant SlSRF protein. The protein was expressed as the *N*-terminus truncated and the *C*-terminus His_6_-tagged form in *E. coli* and purified using a His_6_-tag affinity column. M, protein marker; a, crude protein; b, purified protein. **(B)** SDS-PAGE of the purified SlSRF mutant proteins.

**Supplementary Fig. S8. Stereoselective conversion of 18-oxo-CLA to orobanchol catalyzed by SlSRF.** Incubation with SlSRF and 18-oxo-CLA produces orobanchol selectively. Error bars represent standard deviation from the mean (n = 3 replicates). Asterisk indicates a significant difference between heat-denatured SlSRF and SlSRF (\*\*\**P* < 0.001 by the Student’s *t* test; ns, non-significant).

**Supplementary Fig. S9. *In vitro* enzyme assay of cowpea VuSRF. (A)** SDS-PAGE of the recombinant proteins. Vigun03g413800 and Vigun03g413900 proteins expressed as the *N*-terminus truncated and the *C*-terminus His_6_-tagged forms in *E. coli*. M, protein marker; a, empty vector; b, crude protein of Vigun03g413800; c, crude protein of Vigun03g413900. **(B)** MRM chromatograms of reaction mixtures of recombinant proteins with 18-oxo-CLA. Vigun03g413900 functions as an SRF catalyzing the stereoselective conversion of 18-oxo-CLA to orobanchol.

**Supplementary Fig. S10. Two cyclization pathways from protonated 18-oxo-CLA.** 18-Oxo-CLA can be transformed into two different compounds, **(A)** orobanchol and **(B)** *ent*-2′-*epi*-orobanchol, depending on the value of the dihedral angle *φ* before the cyclization reaction. Left, middle, and right panels show the reactant, transition state, and product structures, respectively. The structures were optimized in vacuo at the level of B3LYP/6-31++G(d,p) theory using Gaussian 16 Rev. C01.

**Supplementary Fig. S11. Relative energy profiles of orobanchol and *ent*-2’-*epi*-orobanchol.** The reactant of the deprotonated compounds (orobanchol/*ent*-2’-*epi*-orobanchol) is 18-oxo-CLA (dashed line). Both orobanchol and *ent*-2’-*epi*-orobanchol are converted from protonated 18-oxo-CLA (solid line).

**Supplementary Fig. S12. *In vitro* enzyme activities of the SlSRF mutants.** MRM chromatograms of reaction mixtures of WT or respective SlSRF mutant with 18-oxo-CLA as a substrate are shown. The signal intensity of each chromatogram is 6.3 ×10^5^.

**Supplementary Fig. S13. D37E side chain model of SlSRF.** The side chain of D37E (deep green) was modeled by ‘mutagenesis’ function implemented in PyMOL 2.5.0. The protein model shown in light green is identical to the reactant structure shown in Fig. 6A.

**Supplementary Table S1.** Distance restraints used for MD simulation models.

**Supplementary Table S2.** MRM transitions used for the detection of SLs with LC–MS/MS.

**Supplementary Table S3.** CDS sequences of synthesized SlCYP722C gene.

**Supplementary Table S4.** Primers used for this study.

**Supplementary Data S1.** List of the top 100 genes with high module membership in the tan co-expression module in cowpea roots.

**Supplementary File S1.** Pymol session file containing the key structures of the BC-ring closure reaction catalyzed by SlSRF.

**Supplementary Movie S1.** A 100-ns MD simulations for the 18-oxo-CLA in SlSRF.

**Supplementary Movie S2.** A 100-ns MD simulations for the 18-oxo-CLA in water solvent.

