## Supplementary Materials for "Insights into stereoselective ring formation in canonical strigolactone: Discovery of a dirigent domain-containing enzyme catalyzing orobanchol synthesis"

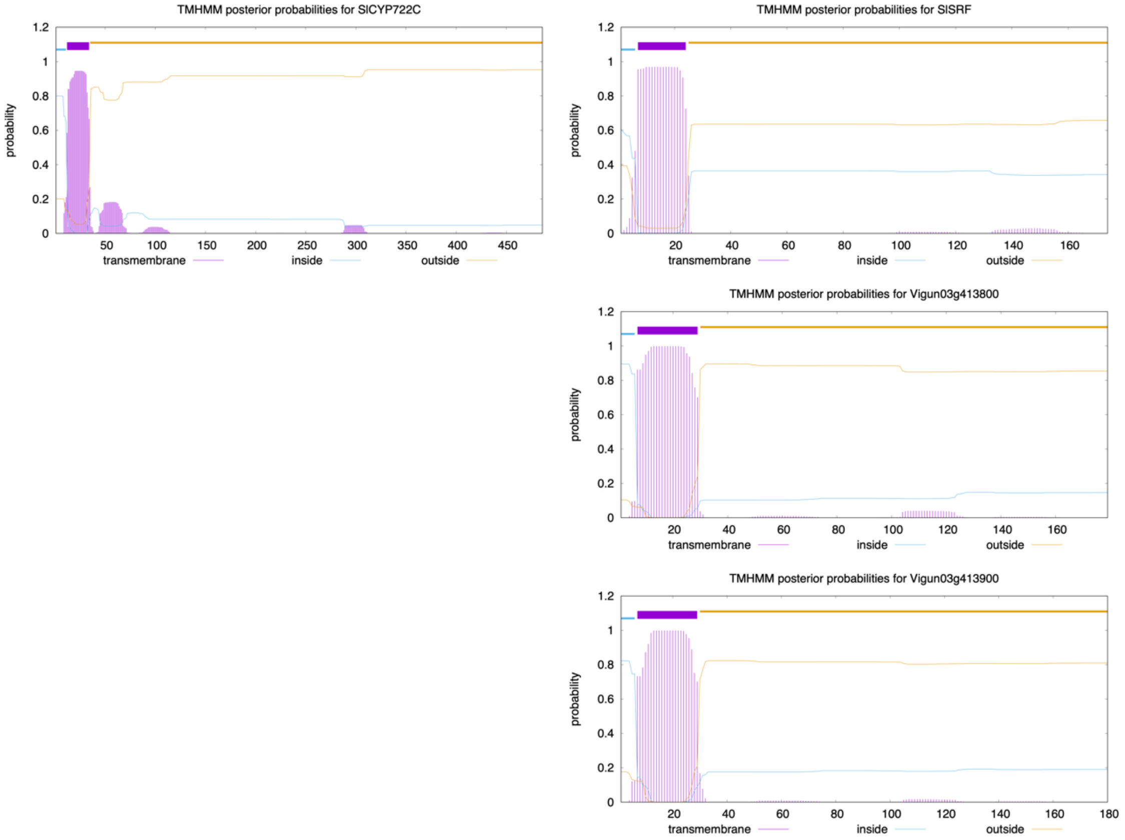


**Supplementary Fig. S1. *In silico* analysis of the SlCYP722C and SRF peptides.** The plot shows the posterior probabilities of inside, outside, and transmembrane helix for each amino acid residue.

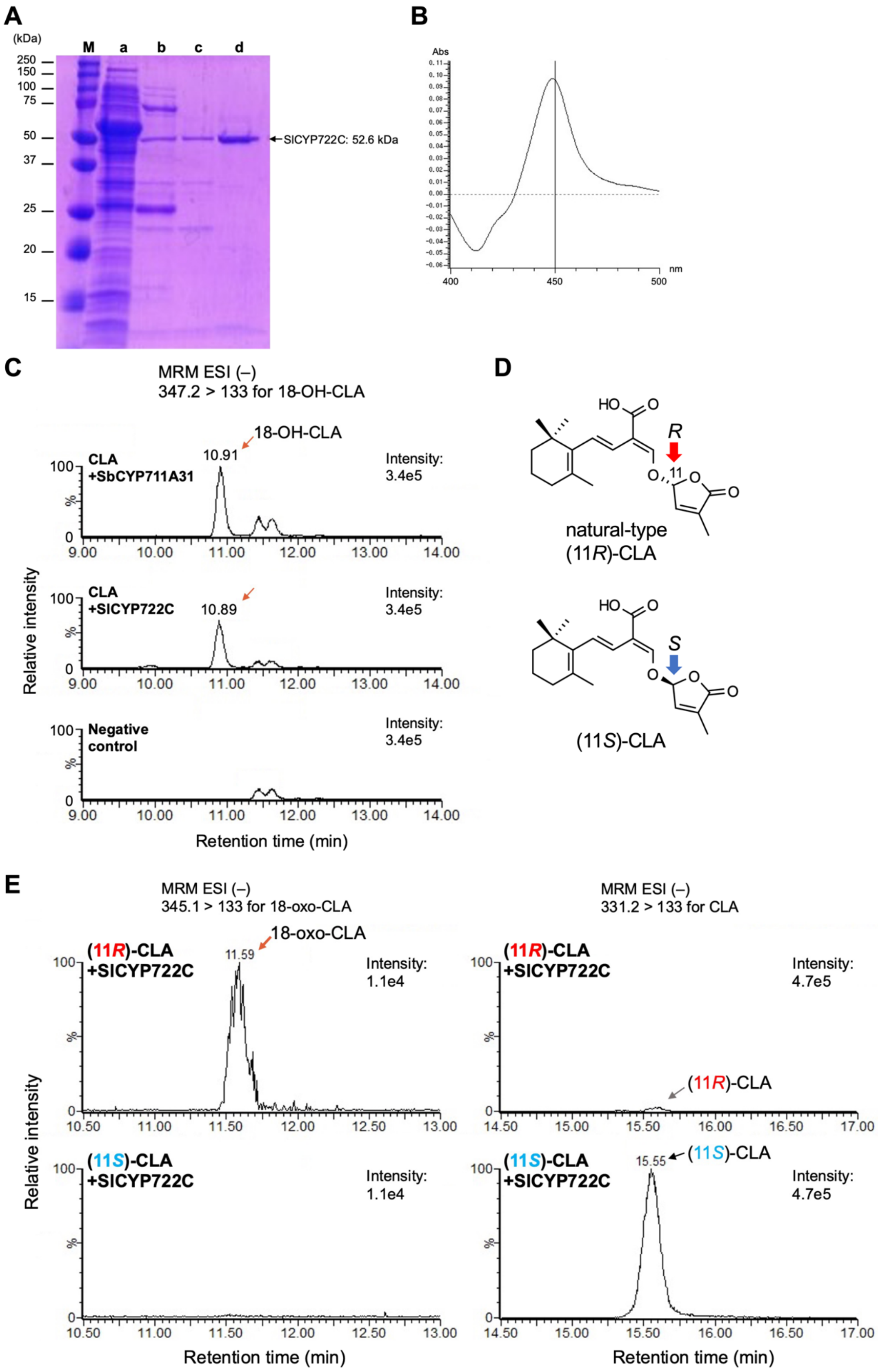
A
(kDa)
M a b c d
250
150
100
75
50
37
25
20
15
SlCYP722C: 52.6 kDa
B
Abs
nm
400
450
500
C
MRM ESI (–)
347.2 > 133 for 18-OH-CLA
18-OH-CLA
10.91
CLA
+SbCYP711A31
Intensity:
3.4e5
CLA
+SlCYP722C
10.89
Intensity:
3.4e5
Negative
control
Intensity:
3.4e5
Relative intensity
Retention time (min)
D
natural-type
(11R)-CLA
(11S)-CLA
E
MRM ESI (–)
345.1 > 133 for 18-oxo-CLA
18-oxo-CLA
11.59
(11R)-CLA
+SlCYP722C
Intensity:
1.1e4
(11S)-CLA
+SlCYP722C
Intensity:
1.1e4
MRM ESI (–)
331.2 > 133 for CLA
(11R)-CLA
+SlCYP722C
Intensity:
4.7e5
(11R)-CLA
(11S)-CLA
+SlCYP722C
15.55
(11S)-CLA
Intensity:
4.7e5
Relative intensity
Retention time (min)

**Supplementary Fig. S2. Heterologous expression of the *N*-terminus truncated SlCYP722C and its activity. (A)** SDS-PAGE of the recombinant SlCYP722C protein. The protein was expressed as the *N*-terminus truncated and the *C*-terminus $His_6$-tagged form in *E. coli* and purified using a series of affinity columns. M, protein marker; a, crude protein; b, purification after $His_6$-tag affinity chromatography; c, purification after DEAE Sepharose anion exchange chromatography; d, final purified protein after CM Sepharose cation exchange chromatography. **(B)** Carbon monoxide **(**CO)-difference spectra of the recombinant SlCYP722C protein. **(C)** Identification of 18-hydroxy-CLA (18-OH-CLA) as the SlCYP722C product. The reaction product of SlCYP722C with 16 Da larger molecular mass than CLA was identified as 18-OH-CLA by comparing the reaction product of SlCYP722C with that of sorghum (*Sorghum bicolor*) SbCYP711A31, which is known to catalyze the conversion of CLA to 18-OH-CLA[57]. **(D)** Structures of (11*R*)-CLA and (11*S*)-CLA. **(E)** Substrate specificity of SlCYP722C. In the case of (11*R*)-CLA as a substrate, 18-oxo-CLA was formed with the decrease of substrate, whereas the reaction did not proceed with (11*S*)-CLA.

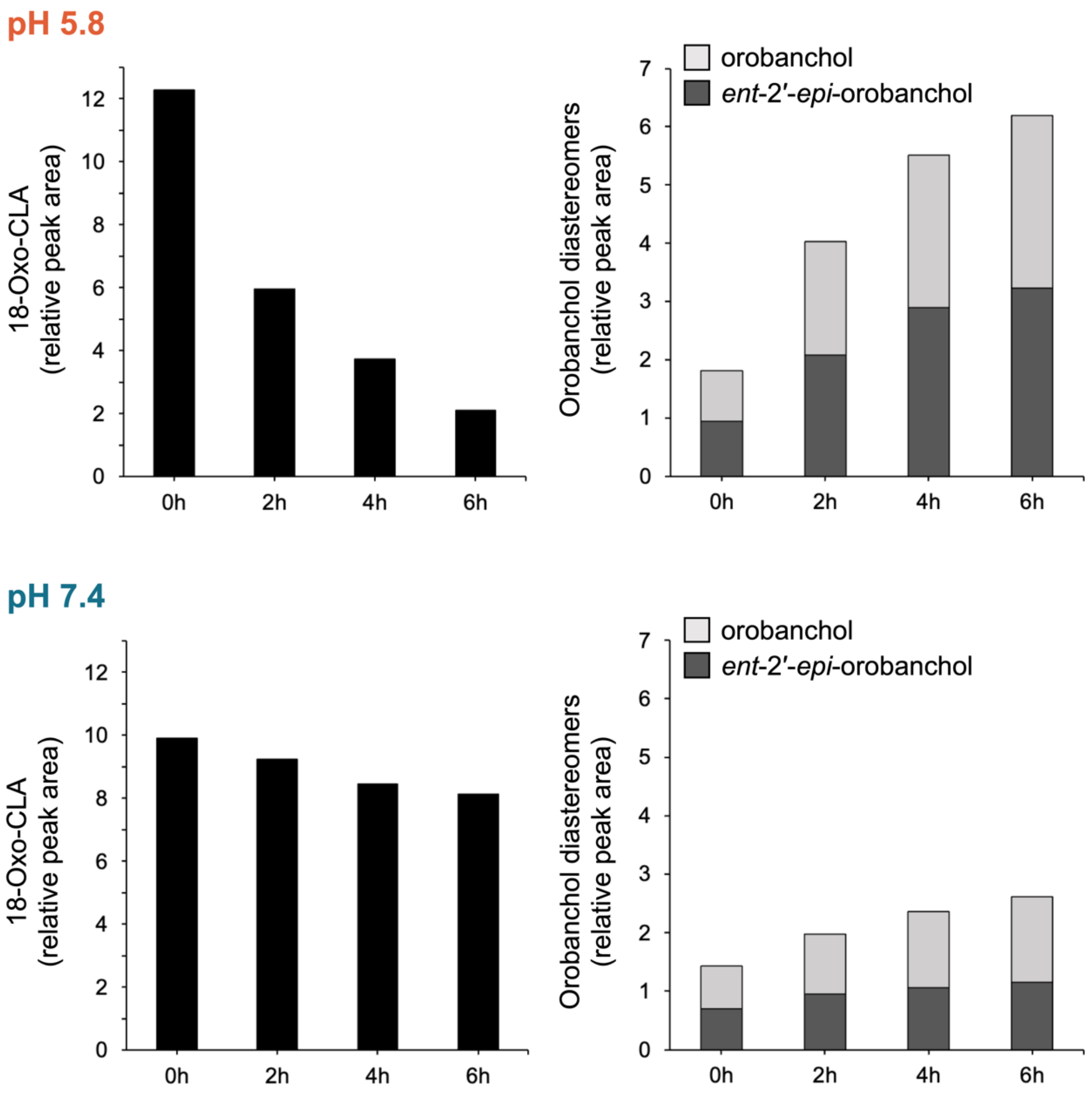


**Supplementary Fig. S3. Formation of orobanchol diastereomers by spontaneous cyclization of 18-oxo-CLA in buffer solution.** The relative amounts of 18-oxo-CLA and orobanchol diastereomers to the internal standard over time differ in sodium potassium buffer at different pH. The decrease in 18-oxo-CLA and the increase in orobanchol diastereomers are more remarkable under acidic conditions.

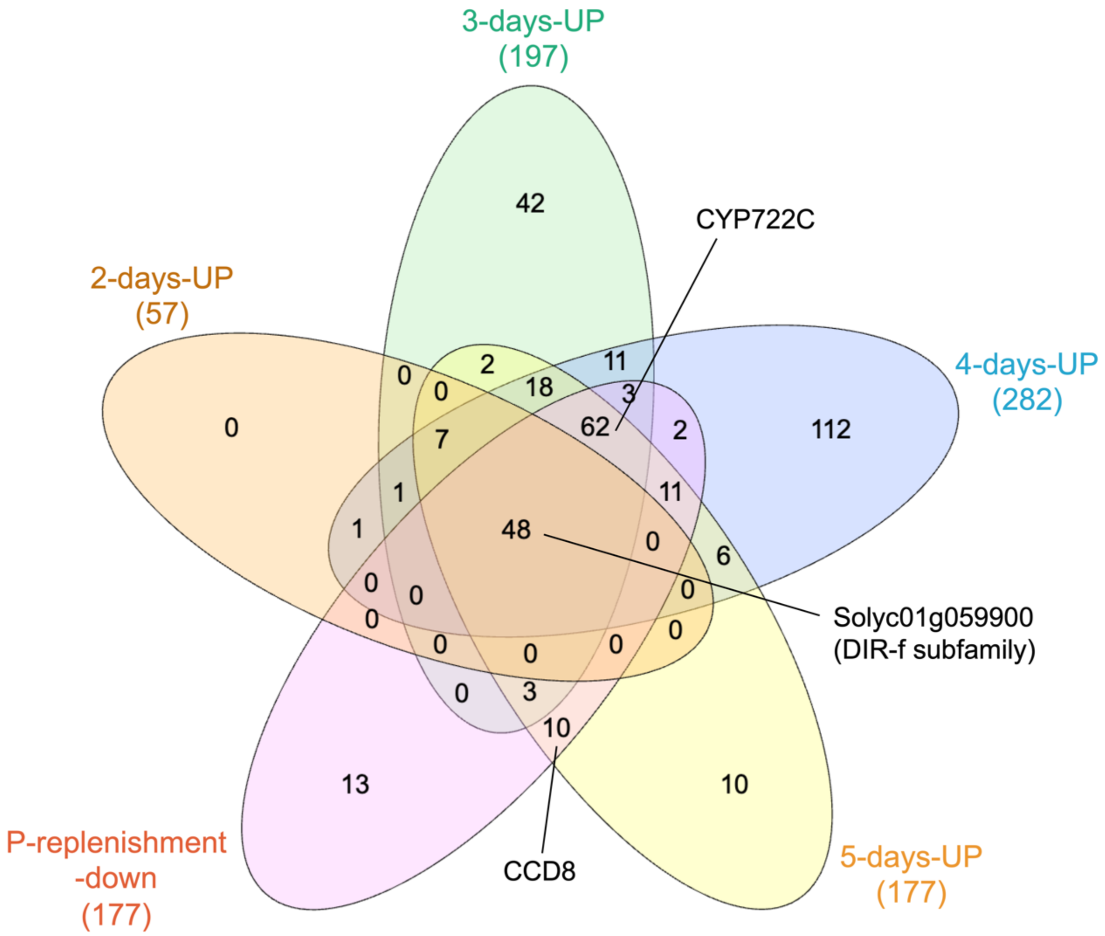


**Supplementary Fig. S4. 48 Differentially expressed genes at the core of the phosphate response in tomato.** Figure adapted and partially modified from Wang et al[29] under the terms of a Creative Commons Attribution 4.0 International License (CC BY 4.0). Venn diagram of up-regulated differentially expressed genes (DEGs) at different time points of phosphate deficiency and down-regulated DEGs by phosphate replenishment.

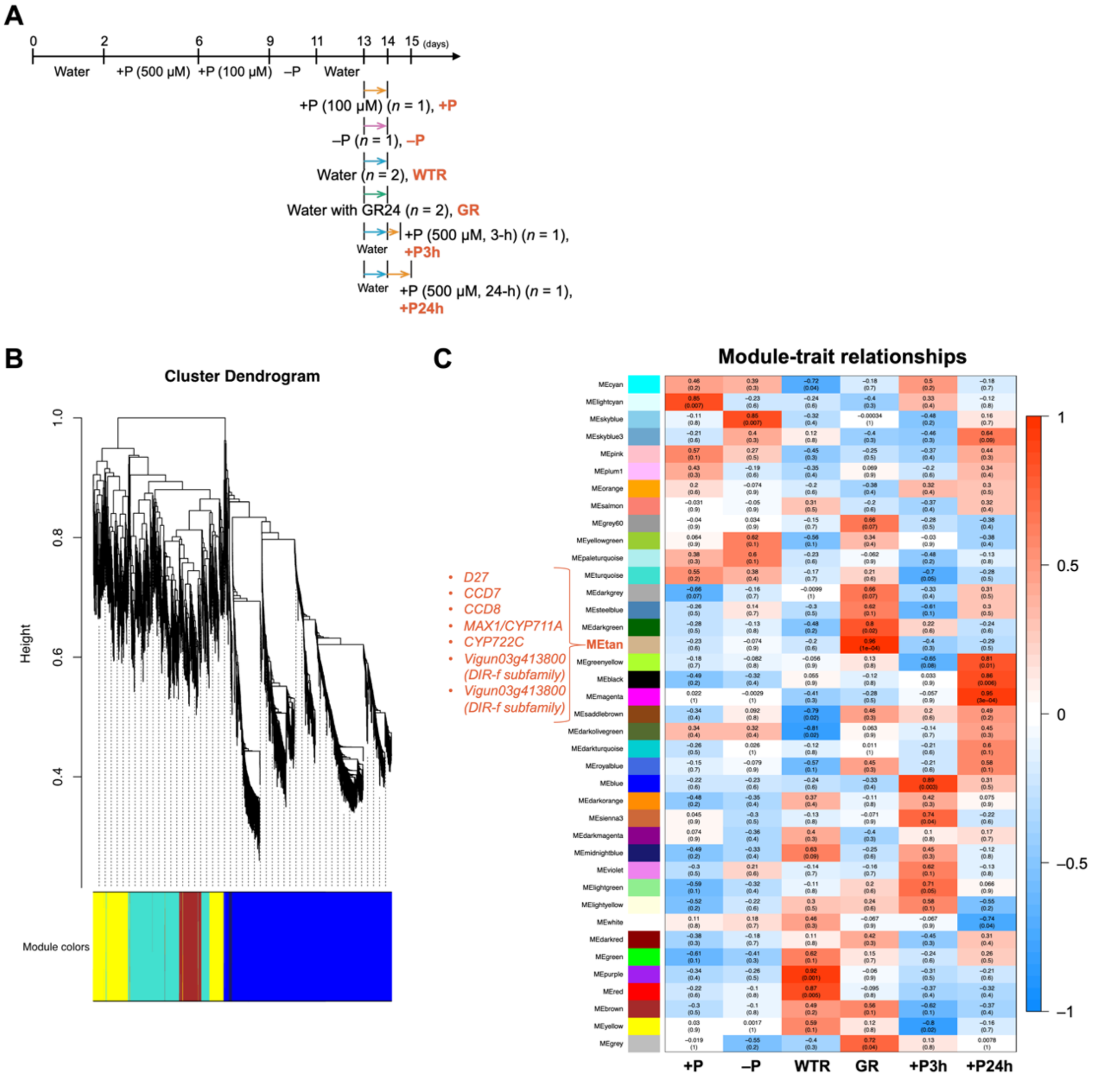


**Supplementary Fig. S5. WGCNA co-expression network and module-trait correlation analysis. (A)** Hydroponic culture conditions of cowpea. Figure adapted and partially modified from our previous paper[13] under the terms of a Creative Commons Attribution NonCommercial Licenses 4.0 (CC BY-NC 4.0). **(B)** Dendrogram plot with color annotation. **(C)** Correlations of WGCNA modules with hydroponic culture conditions (traits) and corresponding P-values. Each row corresponds to each of the 39 co-expression modules and the columns correspond to six traits. The color of each cell indicates the correlation coefficient between the module and traits; the color scale on the right shows module-trait correlation from –1 (blue) to 1 (red).

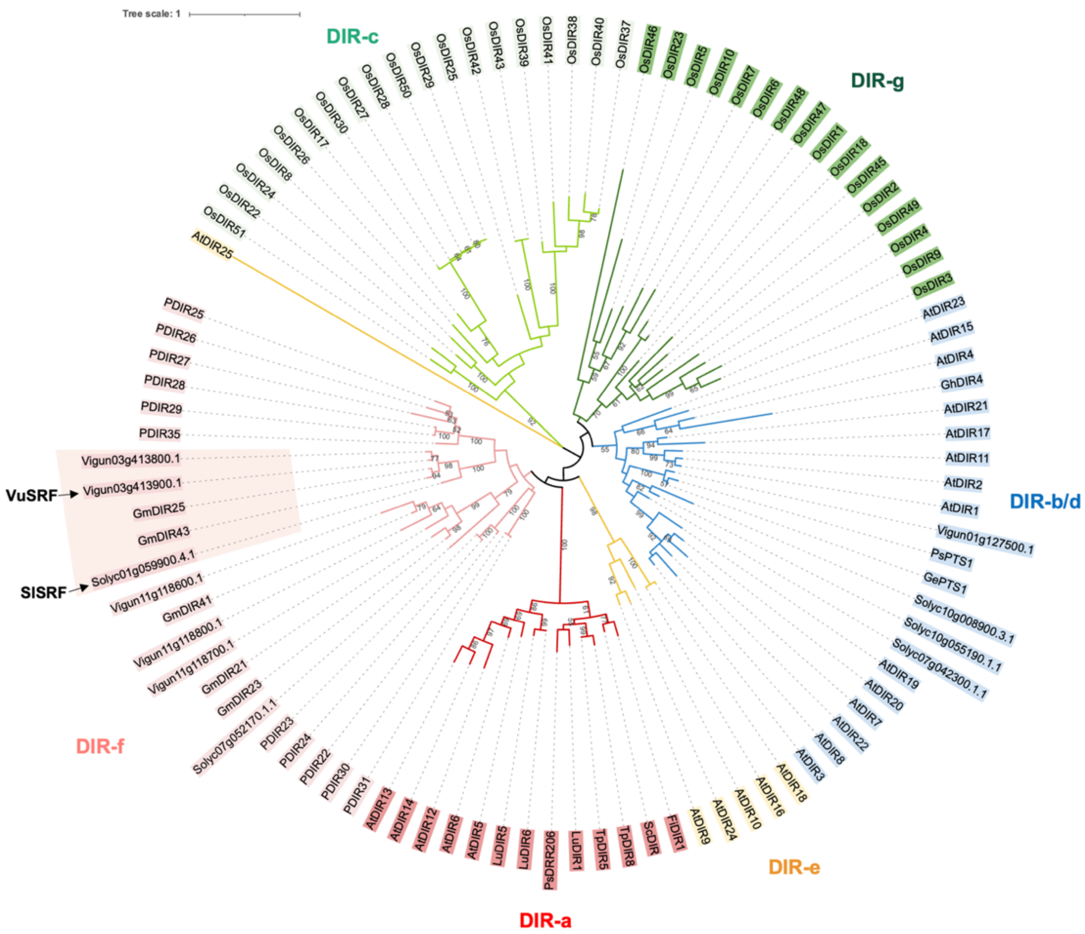


**Supplementary Fig. S6. Phylogenetic tree of DIR sequences.** The amino acid sequences of tomato SlSRF (Solyc01g059900.4.1), cowpea VuSRF (Vigun03g413900), and their paralogs and DIRs from other species were used to conduct the phylogenetic analysis in MEGA X using the maximum likelihood method. DIRs were classified into six groups, DIR-a to DIR-f, based on their phylogenetic relationships with reference to previous report[33] and are indicated by different colors in the phylogenetic tree. The clade containing the SRFs is shaded in pink. At, *Arabidopsis thaliana*; Fi, *Forsythia intermedia*; Ge, *Glycyrrhiza echinata*; Gh, *Gossypium hirsutum*; Gm, *Glycine max*; Lu, *Linum usitatissimum*; Os, *Oryza sativa*; P, *Picea glauca*; Ps, *Pisum sativum*; Sc, *Schizandra chinensis*; Tp, *Thuja plicata*.

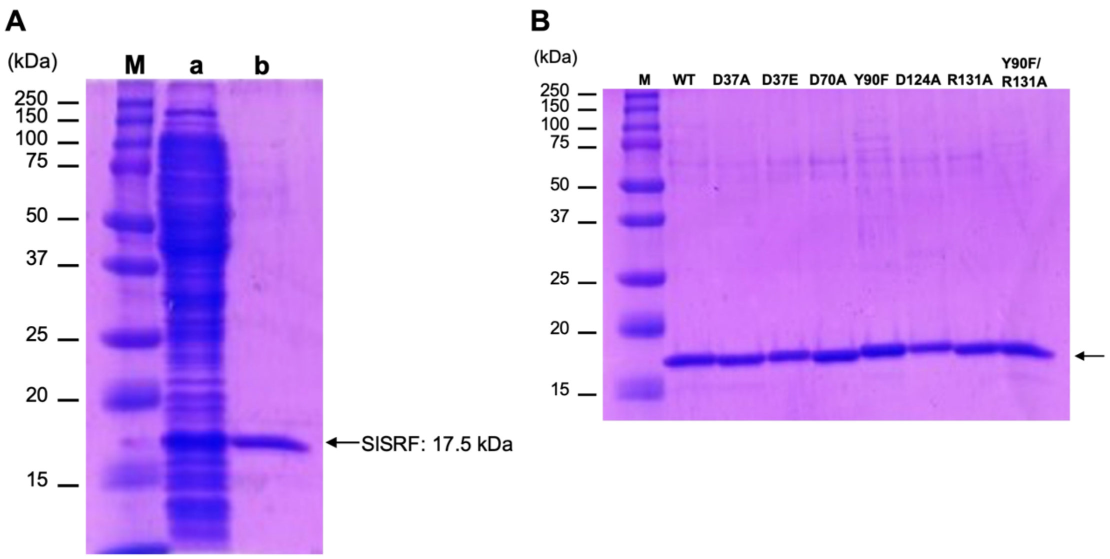


**Supplementary Fig. S7. Purification of recombinant SlSRF protein. (A)** SDS-PAGE of the recombinant SlSRF protein. The protein was expressed as the *N*-terminus truncated and the *C*-terminus $His_6$-tagged form in *E. coli* and purified using a $His_6$-tag affinity column. M, protein marker; a, crude protein; b, purified protein. **(B)** SDS-PAGE of the purified SlSRF mutant proteins.

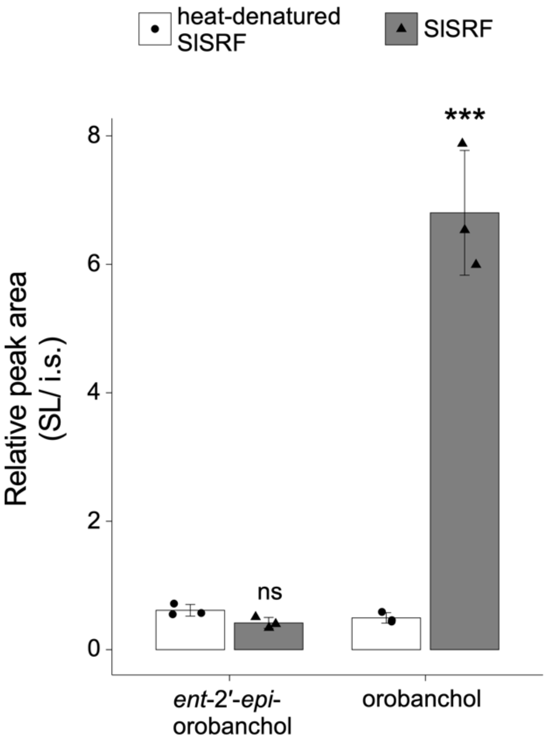


**Supplementary Fig. S8. Stereoselective conversion of 18-oxo-CLA to orobanchol catalyzed by SlSRF.** Incubation with SlSRF and 18-oxo-CLA produces orobanchol selectively. Error bars represent standard deviation from the mean (n = 3 replicates). Asterisk indicates a significant difference between heat-denatured SlSRF and SlSRF (***$P < 0.001$ by the Student's $t$ test; ns, non-significant).

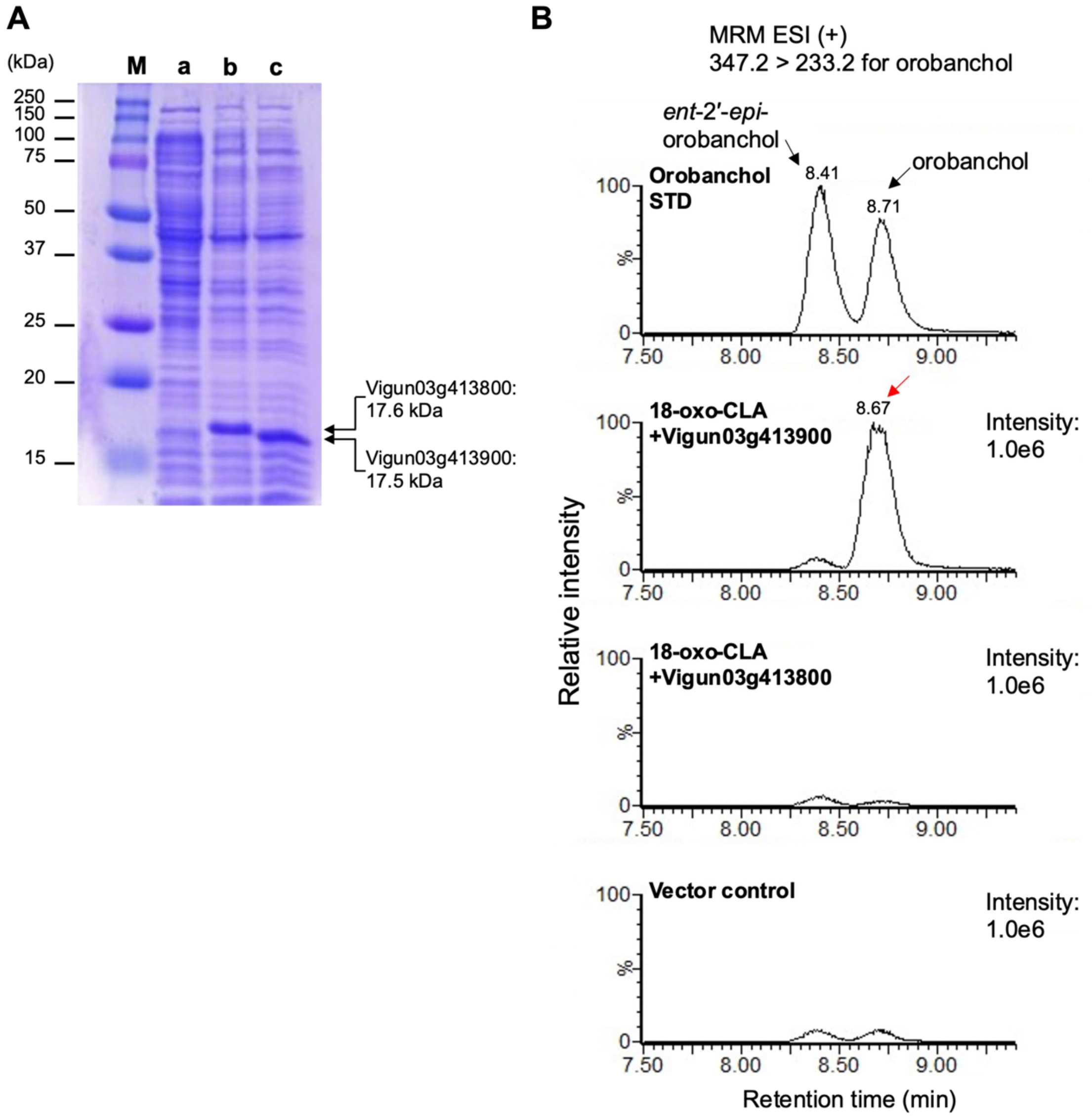


**Supplementary Fig. S9. *In vitro* enzyme assay of cowpea VuSRF. (A)** SDS-PAGE of the recombinant proteins. Vigun03g413800 and Vigun03g413900 proteins expressed as the *N*-terminus truncated and the *C*-terminus $His_6$-tagged forms in *E. coli*. M, protein marker; a, empty vector; b, crude protein of Vigun03g413800; c, crude protein of Vigun03g413900. **(B)** MRM chromatograms of reaction mixtures of recombinant proteins with 18-oxo-CLA. Vigun03g413900 functions as an SRF catalyzing the stereoselective conversion of 18-oxo-CLA to orobanchol.

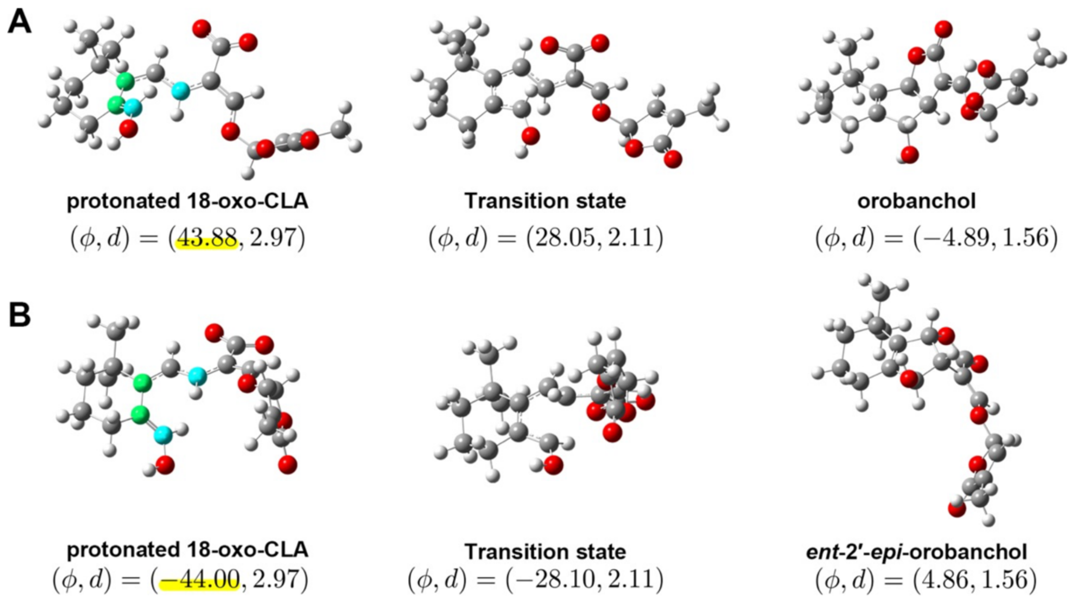


**Supplementary Fig. S10. Two cyclization pathways from protonated 18-oxo-CLA.** 18-Oxo-CLA can be transformed into two different compounds, **(A)** orobanchol and **(B)** *ent*-2′-*epi*-orobanchol, depending on the value of the dihedral angle $\varphi$ before the cyclization reaction. Left, middle, and right panels show the reactant, transition state, and product structures, respectively. The structures were optimized in vacuo at the level of B3LYP/6-31++G(d,p) theory using Gaussian 16 Rev. C01.

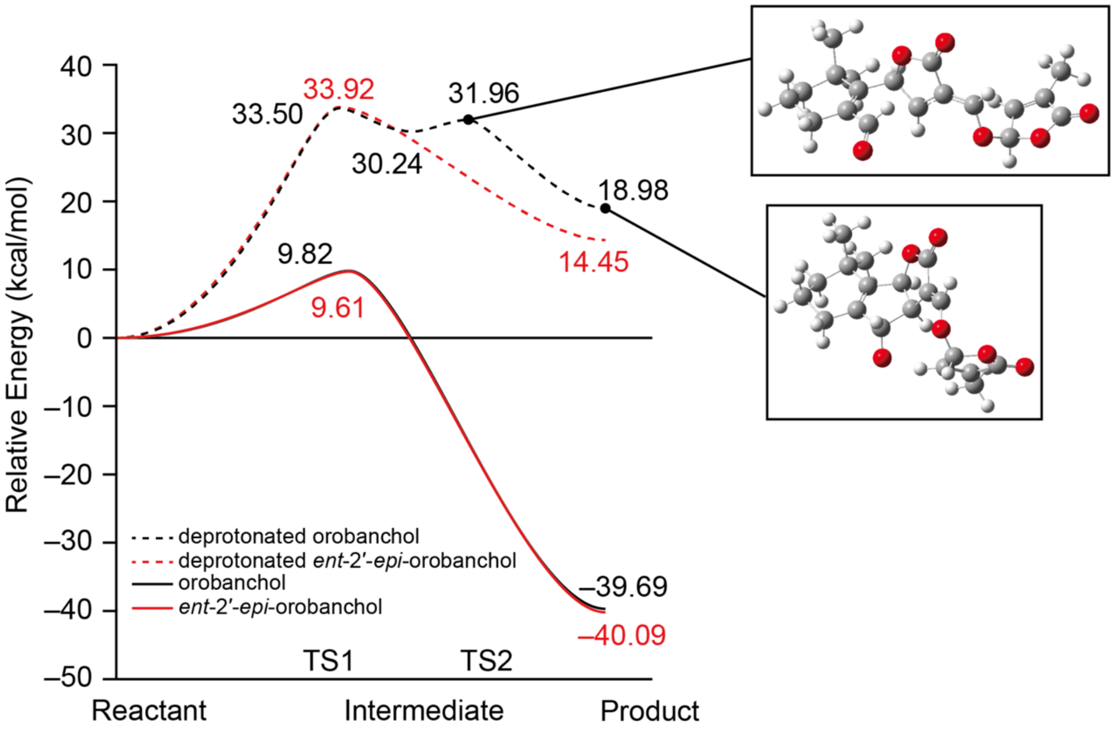


**Supplementary Fig. S11. Relative energy profiles of orobanchol and *ent*-2'-*epi*-orobanchol.**
The reactant of the deprotonated compounds (orobanchol/*ent*-2'-*epi*-orobanchol) is 18-oxo-CLA (dashed line). Both orobanchol and *ent*-2'-*epi*-orobanchol are converted from protonated 18-oxo-CLA (solid line).

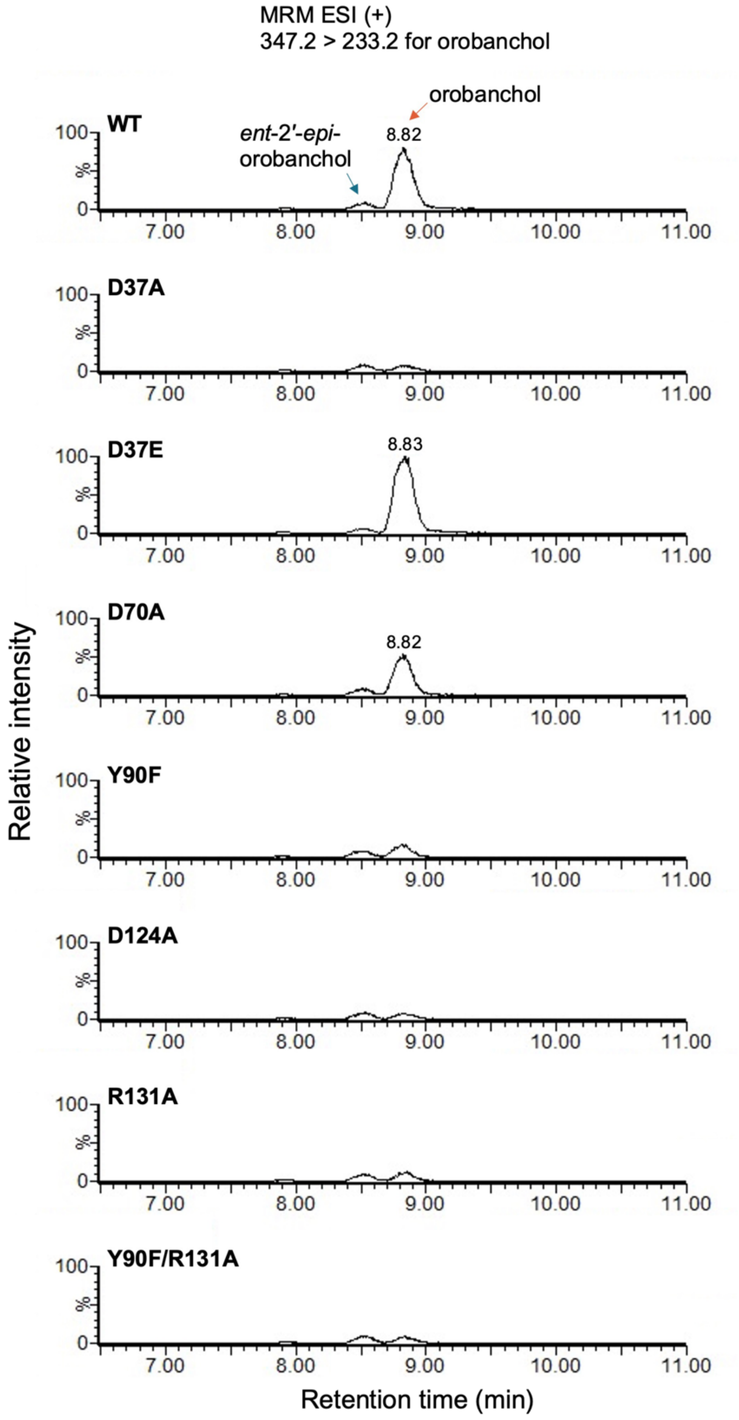


**Supplementary Fig. S12. *In vitro* enzyme activities of the SlSRF mutants.** MRM chromatograms of reaction mixtures of WT or respective SlSRF mutant with 18-oxo-CLA as a substrate are shown. The signal intensity of each chromatogram is $6.3 \times 10^5$.

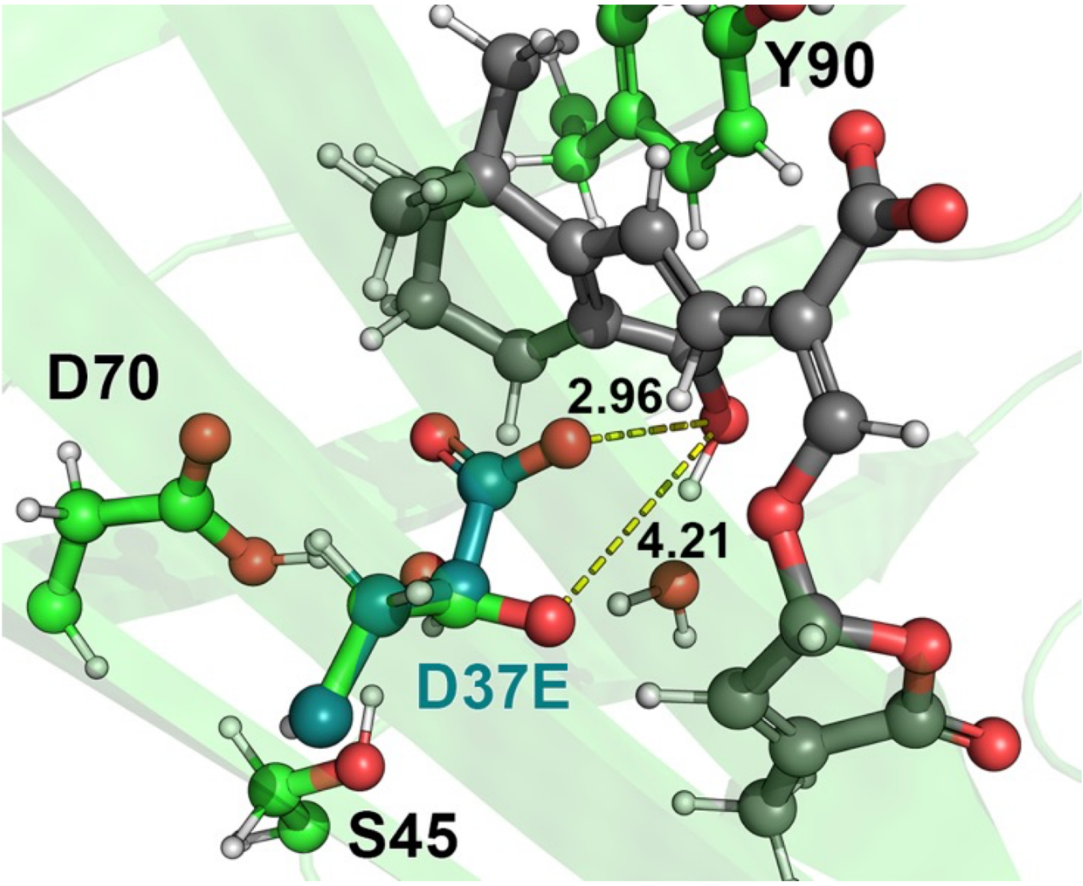


**Supplementary Fig. S13. D37E side chain model of SlSRF.** The side chain of D37E (deep green) was modeled by 'mutagenesis' function implemented in PyMOL 2.5.0. The protein model shown in light green is identical to the reactant structure shown in Fig. 6A.

**Supplementary Table S1.**

Distance restraints used for MD simulation models.

| Atom 1 | Atom 2 | Equilibration distance *r* (Å) | Force constants *k* (kcal mol$^{-1}$ Å$^{-2}$) |
|---|---|---|---|
| protonated Asp37–Oδ2 | 18-oxo-CLA–aldehyde group | 2.90 | 2.0 |
| Asp124–Cγ | Arg131–Cζ | 4.00 | 2.0 |
| Arg131–Nη | 18-oxo-CLA–carboxy group | 3.00 | 2.0 |
| 18-oxo-CLA–C11 | 18-oxo-CLA–C9 | 3.20 | 2.0 |

**Supplementary Table S2.**

MRM transitions used for the detection of SLs with LC–MS/MS.

| SL | ESI mode | Transition (*m/z*) | Cone voltage (V) | Collision energy (eV) |
|---|---|---|---|---|
| orobanchol isomers | + | 347.2 > 233.2 | 32 | 14 |
| 18-oxo-MeCLA | + | 361.2 > 97.0 | 35 | 20 |
| CLA | − | 331.2 > 113.0 | 30 | 18 |
| 18-oxo-CLA | − | 345.1 > 113.0 | 30 | 18 |
| 18-OH-CLA | − | 347.2 > 113.0 | 30 | 18 |

**Supplementary Table S3.**

CDS sequences of synthesized SlCYP722C gene.

| Gene name | Sequence (5′–3′) |
|---|---|
| ΔN34-SlCYP722C-His$_6$ | ATGGCCTCGAAAGGGGCGAAAAACGCCAAATCATGCATTCCGGGTAGTCT<br>GGGCATACCGTTTGTTGGCGAAACCTTCGCACTGTTATCCGCTACCAACA<br>GCGTCAAGGGCTGCTATGAATTTGTCAGACTGCGGCGTGAACGGCATGGC<br>AAGTGGTTCAAAACCCGCATATTCGGAAAGATTCACGTCTTTGTTCCGTC<br>AGTGGAAGGTGCGAAAGCCATTTTCACCAACGATTTTGCGCTCTTCAACA<br>AAGGCTACGTCAAAAGCATGGCAGATGCGGTTGGCAAAAAGTCTCTGCTC<br>TGTGTGCCGCAAGAATCGCATAAACGCATTCGTCGCCTGTTAAGTGATCC<br>GTTTAGCATGAACAGTCTGTCGAAATTTGTACAACGTTTTGACGAGATGC<br>TGTATGAACGCTTGAAGAAAGTTCAGAAACAACGCAAGAGTTTTACCGTG<br>TTAGATTTCAACATGAAAACGACGTTTGATGCGATGTGCGACATGTTGAT<br>GAGCATCAAGGACTCGTCTGTCGTTGAACAGATTGAGAAAGATTGCACTG<br>CTATCAGCGATGCAATGCTGTCTTTTCCGGTTATGATTCCGGGAACGCGCT<br>ATTTCAAGGGCATCAAAGCACGTGGTAGGCTGATGGAAACCTTCAAAGG<br>GATGATTGCGGCACGTCGTAATGGGAAAGAGTACTACGAAGATTTCCTGC<br>AGAGCATGCTGGAAAAAGACAGTTGTCCAGCCAATCAGAAACTGGACGA<br>TGAAGAGATCATGGATAACCTGCTTACTTTGATCATCGCAGGTCAAACCA<br>CTACAGCTGCGGCGATGATGTGGTCCGTGAAATTTCTGGATGAGCACCGT<br>GAAGCCCAGAATCGCTTACGAGAAGAGCAGTTGAGCATCTTACGTTCCAA<br>ACCAACAGGTGCCTTACTGACGATGGATGACCTGAACTCCATGTCTTATG<br>CCAGCAAAGTGGTGAAAGAAACGCTGCGGATGAGCAATGTGCTCCTGTG<br>GTTTCCACGAGTAGCTCTAAATGACTGCTCGATTGAGGGTTTTGAGATCA<br>AAAAAGGCTGGCATGCGAATATCGATGCGACTTGCATCCACTATGATCCT<br>ACGCTGTACAAAGACCCTATGCAGTTTAATCCGTCACGTTTCGACGAAAT<br>TCAGAAACCGTATTCTTATATTCCCTTTGGAAGCGGTCCACGTACCTGTTT<br>GGGCATTAACATGGCCAAAGTGACCATGCTGGTGTTCCTTCATCGCCTGA<br>CCACCGGTTACAAGTGGACAGTAGACGATCCCGATCGCTCCCTAGAACGC<br>AAAGCGCATATTCCTCGCCTTCGTTCAGGCTGTCCGATTACACTTACGGCT<br>CTCCCGGATGAACATCACCATCACCACCATTAG |

**Supplementary Table S4.**

Primers used for this study.

| Primer name | Primer sequence (5′–3′) | Note |
|---|---|---|
| **Cloning for protein expression** | | |
| ΔSP-SlSRF-His-F | gccgcat**atg**AACTTTAAAAAACAAA | Gene specific sequences are capitalized. Start and Stop codons are highlighted in bold. Restriction enzyme sites are underlined. |
| ΔSP-SlSRF-His-R | cggcggtaccATAGTGCTTAACAGTAACATCA | |
| ΔSP-Vigun03g413800-His-F | gccgcat**atg**CAACCAAAGCAAACCAATCTT | |
| ΔSP-Vigun03g413800-His-R | cggcggtaccGTAATGTTTGAGTGTTACATT | |
| ΔSP-Vigun03g413900-His-F | gccgcat**atg**CAACCCAAGCAAACCAATCTC | |
| ΔSP-Vigun03g413900-His-R | cggcggtaccGTAATGTTTGAGTGTTACAGT | |
| **PCR based mutagenesis** | | |
| SlSRF-D37A-F | GTCCATGCGTATTTAAGTGGACATGACACATCTGC | The underlined codon corresponds to the mutated amino acid. |
| SlSRF-D37A-R | TAAATACGCATGGACATAAAATGCTAGTTTTGTTT | |
| SlSRF-D37E-F | GTCCATGAATATTTAAGTGGACATGACACATCTGC | |
| SlSRF-D37E-R | TAAATATTCATGGACATAAAATGCTAGTTTTGTTT | |
| SlSRF-D70A-F | GTTGATGCGCCAGTAACAGAAGGTCCAGATTTAAA | |
| SlSRF-D70A-R | TACTGGCGCATCAACAGCAATAATTGTACCAAAAG | |
| SlSRF-Y90F-F | GGAATGTTTATAAATTCACAATTGGATGGCAAAGG | |
| SlSRF-Y90F-R | ATTTATAAACATTCCTTGGGCTCTACCAATTAATT | |
| SlSRF-D124A-F | GGTGCTGCGTTATTTGCAATGAAAGAAAGGGAATT | |
| SlSRF-D124A-R | AAATAACGCAGCACCTTGAATCTCCAAAGTACTCC | |
| SlSRF-R131A-F | AAAGAAGCGGAATTTTCAATTGTGTCTGGGACTGG | |
| SlSRF-R131A-R | AAATTCCGCTTCTTTCATTGCAAATAAATCAGCAC | |
